# Separating biological signal from statistical artifact in trait trade-off inference

**DOI:** 10.64898/2026.10.03.756330

**Authors:** Garrison W. Bullard, Donald A. Jackson

## Abstract

Trade-offs are commonly assumed when the statistical parameters used to define biological traits are negatively correlated. However, parameter correlations can emerge from the geometry of the model, rather than underlying biological signal. We characterize this phenomenon generally, and develop a framework to parse biological signal from statistical artifact in parameter correlations. Using the widely reported trade-off between thermal sensitivity and static upper critical thermal limit as a case study, we show that an extreme correlation between these parameters is expected even in the absence of a real trade-off. However, our method reveals that there is, in fact, a biological correlation underneath this spurious artifact. We confirm this finding, as independent genome-wide association analysis reveals genetic variation among trade-off phenotypes, but none corresponding to net thermal tolerance, suggesting that adaptive capacity to a warming climate may be severely limited. This particular trade-off is therefore biologically supported, illustrating how biological signal can be obscured by estimation artifact, thus evading correlation-based detection. We broaden our analysis, showing that artifact is present across biological systems and model types, although its manifestation may vary. Analytical results and simulations reveal how sampling design, model choice and parameterization govern the artifact, providing practical guidance for experimental planning. Our findings identify both a problem and a solution: correlations among model parameters cannot be interpreted biologically without first accounting for the structure imposed by the model and data, but genuine biological relationships can still be recovered when that structure is explicitly modeled.

## Introduction

Biologists routinely conclude that traits trade off when their defining statistical parameters correlate, but that correlation can arise from pure mathematical artifact rather than biological signal (1–4). While some traits can be empirically measured (e.g., height, weight), many are not directly observable and are instead inferred from statistical and mechanistic models. Biological models are assumed to represent a natural system, and their parameters are often interpreted as biological traits (e.g., fitness, IQ, heritability) (5, 6). Correlations among those parameters are therefore often interpreted as relationships between the traits themselves. In particular, when favorable values of one parameter are associated with unfavorable values of another, the traits are frequently said to trade off biologically (7). However, this process of interpretation has been criticized for lacking rigor, as correlations among parameter values do not necessarily represent biological relationships among their corresponding traits when multiple parameters are jointly estimated (2, 8–10).

Even with high-quality data and well-designed models, multiple parameter combinations can often explain the observations with comparable likelihood because changes in one parameter can be offset by changes in another (11). Such sets of parameters are called “non-identifiable” (12, 13). Biological datasets can be particularly vulnerable to non-identifiability, as these systems are inherently variable, while cost and time demands constrain sample sizes (14, 15). Under these conditions, ambiguity in how variation should be attributed among parameters induces structured uncertainty and bias in their estimated values (13). This structure can generate compelling patterns in parameter means, variances, covariances, and correlations even when no corresponding structure exists biologically. Thus, poorly identified parameters cannot be automatically interpreted as traits. Failure to recognize this distinction can yield unfounded inferences about a trait’s biological role. While non-identifiability can alter many kinds of parameter estimates, we approach this problem through the lens of spurious correlations.

Concerns about spurious correlations and their ability to distort trait inferences have been raised by ecologists, but their cause and magnitude of influence are general points of uncertainty and confusion (2, 8–10). Mechanisms of spurious correlation have been proposed and solutions developed for particular cases, such as relating two ratios of the same variable (2), paired observation data (8, 16), or Strehler-Mildvan correlation in the Gompertz model of aging (10). These cases, however, have largely been treated as separate statistical problems rather than as manifestations of a common problem of parameter non-identifiability. Thus, a general framework is needed to determine the parameter correlation expected from the model and data alone and whether the observed pattern exceeds that expectation. Even in the specialized applications where corrective methodology is available, multiple sources of spurious correlation may be present, making “corrected” estimates difficult to interpret. For example, an estimate of correlation that is corrected for artifact A might still be spurious due to artifact B. Additionally, adjusted correlations are typically not corroborated with empirical evidence, leaving uncertainty as to whether any given parameter correlation truly indicates a biological trade-off.

Here, we present a general framework for separating the variance-covariance structure expected from parameter non-identifiability from structure consistent with biological variation. We test this methodology using a frequently reported trade-off between thermal sensitivity and the upper critical temperature. Across ectotherm populations, these parameters exhibit an extreme correlation frequently less than −0.9, apparently providing overwhelming evidence of a biological constraint (17–21). However, both parameters are inferred from the same model over a restricted predictor range and related by a common variable, and we show how under these conditions, poor identifiability may itself produce an extreme correlation. This system therefore provides a stringent test of whether genuine biological trade-offs can be recovered when the apparent evidence for them is dominated by statistical artifact.

Using simulations, we show how finite sampling across a narrow predictor range and nonlinear reparameterization can produce strong parameter correlations in the absence of biological covariation. We then derive the parameter variance/covariance structure expected in the absence of biological covariation. Applying this framework to an empirical dataset of thermal tolerance estimates for 100 isogenic lines of *Drosophila melanogaster* shows that extreme correlation between estimated thermal sensitivity and upper critical temperature is largely expected from non-identifiability alone. Nevertheless, the observed relationship is even more extreme than these null estimates. We corroborate this residual biological signal by pairing dimensionality reduction with genomic data, finding that all detectable genetic variation lies within the thermal tolerance trade-off axis. Thus, our sta-tistical and genomic analyses independently support the existence of the proposed thermal tolerance trade-off, albeit likely at a weaker magnitude than raw parameter correlation would suggest. In contrast, the genomic analysis finds no genetic variation within the orthogonal net thermal tolerance axis, providing striking evidence that the potential for population adaptation to a warming climate is severely limited.

Testing our framework against 25 literature datasets with previously established trade-offs validates that our methodology can distinguish biological signal from spurious correlation. Deploying our test on three more literature “trade-off” datasets using varied linear mixed models shows that artifactual influences on parameter correlational direction and magnitude are commonly strong across biological fields, with biases differing strongly by modeling technique. We discuss how this methodology can extend to non-linear systems.

We conclude by quantitatively determining the empirical and analytical sources of variance/covariance bias, enabling researchers to incorporate identifiability theory into experimental design. As such, our findings provide both a warning and a path forward: correlations among model parameters cannot be interpreted biologically without first accounting for the structure imposed by the model, data, and sampling design, but genuine biological relationships can still be recovered when that structure is explicitly modeled or genetically substantiated.

## Exploration and results

### Sampling error combined with reestimation induces altered correlation structure

Empirical biology frequently involves relating some response variable to one or more predictor variables. Scientists take measurements of the response at a finite number of predictor values and fit models to estimate “parameter-traits” that characterize the resulting biological relationship. The most common of these relationships is the linear model. We generated a two-dimensional linear relationship (Figure 1A), where the orange line is the “true” data-generating relationship (representing the underlying statistical population), and each black dot might represent a single observation obtained in the lab or field. In practice, the generating relationship is unknown, but researchers aim to estimate it using the sampled data. Datasets are typically limited to a narrow predictor range (shaded region in Figure 1B), within which researchers would typically fit an ordinary least-squares regression. This estimated line would approximate (but not equal) the generating distribution (dark pink line, Figure 1B). However, there would be some level of non-identifiability in this linear estimation–many candidate parameter combinations may yield near-identical fits (light pink lines, Figure 1B). Note that all of these candidate lines spread out greatly from a pivot point at the data centroid–a defining characteristic of poor identifiability in linear regression (22). Candidate lines with steeper (more negative) slopes must have higher y-intercepts and lower x-intercepts, so the slope and intercepts of candidate lines are tightly correlated due to model uncertainty. If we simulate data and reestimate the line of best fit many times (as in Figure 1B), we obtain spuriously correlated parameter estimates, with some sampling noise (Figure 1C, D). Note that the variance of each parameter is artificially inflated, resulting in high estimation error for individual observations (Figure 1C, D). While we demonstrate this effect with two dimensions, the same problem emerges (often with higher frequency) in multidimensional systems–in fact, its absence is rare in complex linear and non-linear models (11, 23). Next, we see how this mechanism plays out in practice, using thermal tolerance as a model system.

**Fig. 1.**
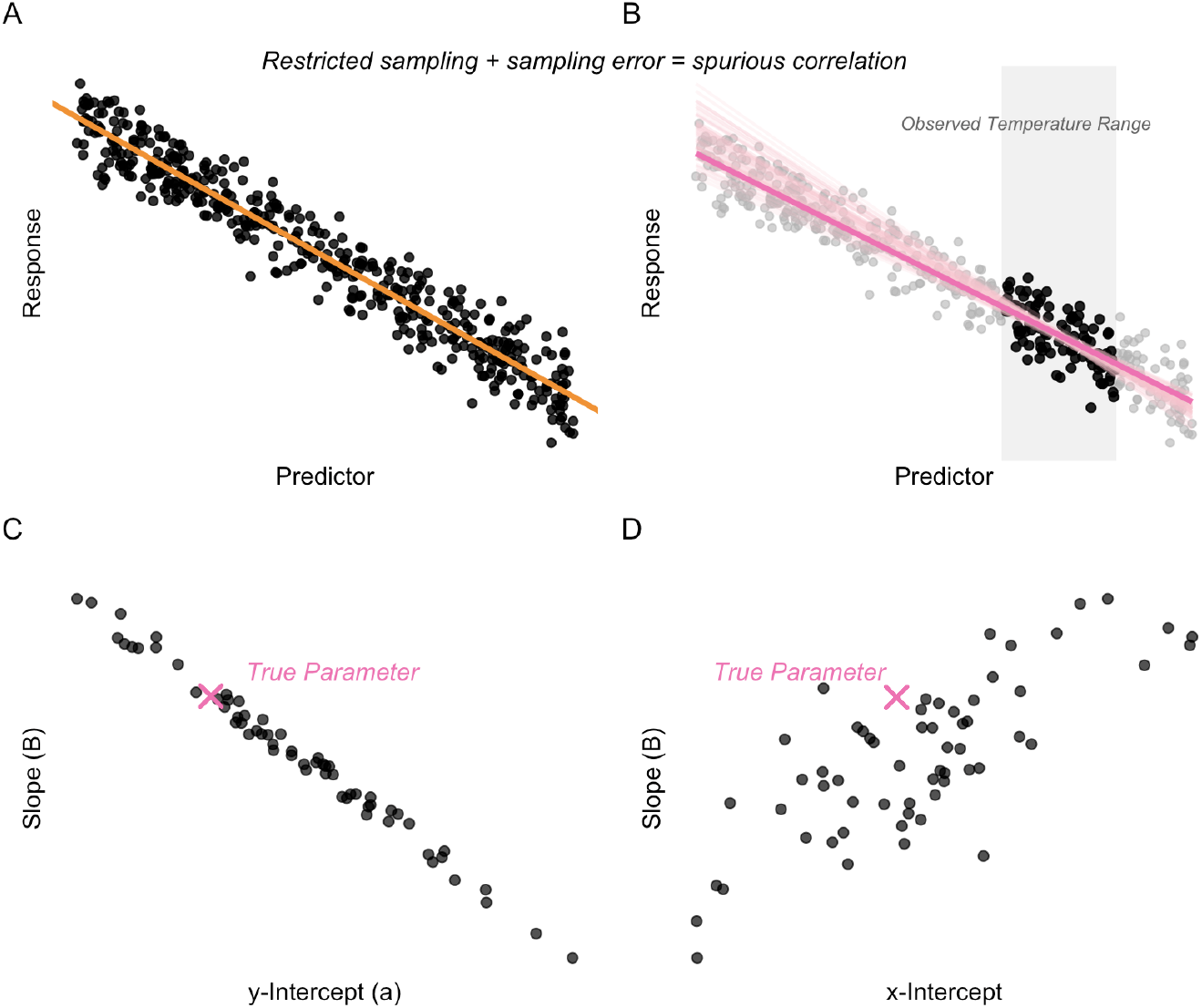
Demonstration of non-identifiability via linear regression. **Panel A:** Many sampled data points (black dots) from a linear generating distribution (orange line). **Panel B:** Estimated least-squares fit (bold pink line) based on data within a restricted predictor range (gray shaded region). Light pink lines denote similarly likely fits obtained by bootstrap sampling the observed data (with replacement) and refitting the line. **Panel C:** Slope/y-intercept pairs for lines from panel B form a spurious correlation. The generating parameter pair is denoted by a pink X. **Panel D:** Slope/x-intercept pairs for lines from panel B form a spurious correlation. The generating parameter pair is denoted by a pink X.

### Artificial correlations in thermal tolerance: non-identifiability + reparameterization

Thermal tolerance in ectotherms is commonly assessed in two ways: dynamic techniques (individuals are subjected to a constant temperature increase, and temperature at failure is recorded) and static techniques (individuals are exposed to a constant temperature, and time at failure is recorded) [failure = an observable endpoint such as death or coma] (24). While each of these approaches condenses thermal tolerance to a single metric (temperature or time), it has long been recognized that ectotherm thermal tolerance is a two-dimensional trait, most accurately described by the relationship between time to failure and temperature (17, 25).

The Thermal Death Time (TDT) framework describes the strong (typically *R*^2^ *>* 0.95) log-linear relationship between a “stressful” temperature (*T*) and an organism’s time to failure at that constant temperature (*t*_*f*_) (1):

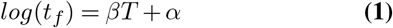

This relationship is typically reparameterized into a more biologically interpretable form, with its parameters explicitly treated as traits, where the static upper critical thermal limit, *sCT*_*max*_ = *− α/β*, is the temperature at which an individual fails in*−* 1 minute, and *z* = −1*/β* is the thermal sensitivity constant (discussed more below) (17).

It is claimed that the optimal thermal generalist “jack of all trades” phenotype would be one that can withstand high temperatures (high *sCT*_*max*_), for a long time (low *z*) (17). However, such a phenotype is deemed theoretically impossible (analogous to a Darwinian Demon), as empirical studies have repeatedly found a strong positive correlation between *z* and *sCT*_*max*_ (commonly >0.9) at several organizational levels, from within populations to between species [e.g., (17, 20)]. This correlation has been deemed the result of a physiological trade-off by the thermal ecology community, although no mechanism for this trade-off has been verified (17, 20, 21). The existence of such a trade-off would theoretically pose a limitation to the adaptation and acclimation of organisms to a warming climate.

Figure 2 demonstrates that the empirically observed correlation between *z* and *sCT*_*max*_ emerges from the same statistical artifact as in Figure 1. Studies that identify a trade-off between *z* and *sCT*_*max*_ also find strong correlations between *α* and *β* (typically*−* 1 *< r < −* 0.9); although this is not usually interpreted physiologically [e.g., (21)]. Figure 2A-C show the artifactual emergence of the slope-intercept correlation under TDT conditions, and how reparameterization to *z/sCT*_*max*_ form flips this correlation to positive, although its strength is weakened by the nonlinear transformations (Figure 2D). The correlation flip caused by reparameterization (Figure 2C vs 2D) can be compared to that of Figure 1 (Figure 1C vs 1D), where *sCT*_*max*_ is the x-intercept and *z* is positively related to *β*.

**Fig. 2.**
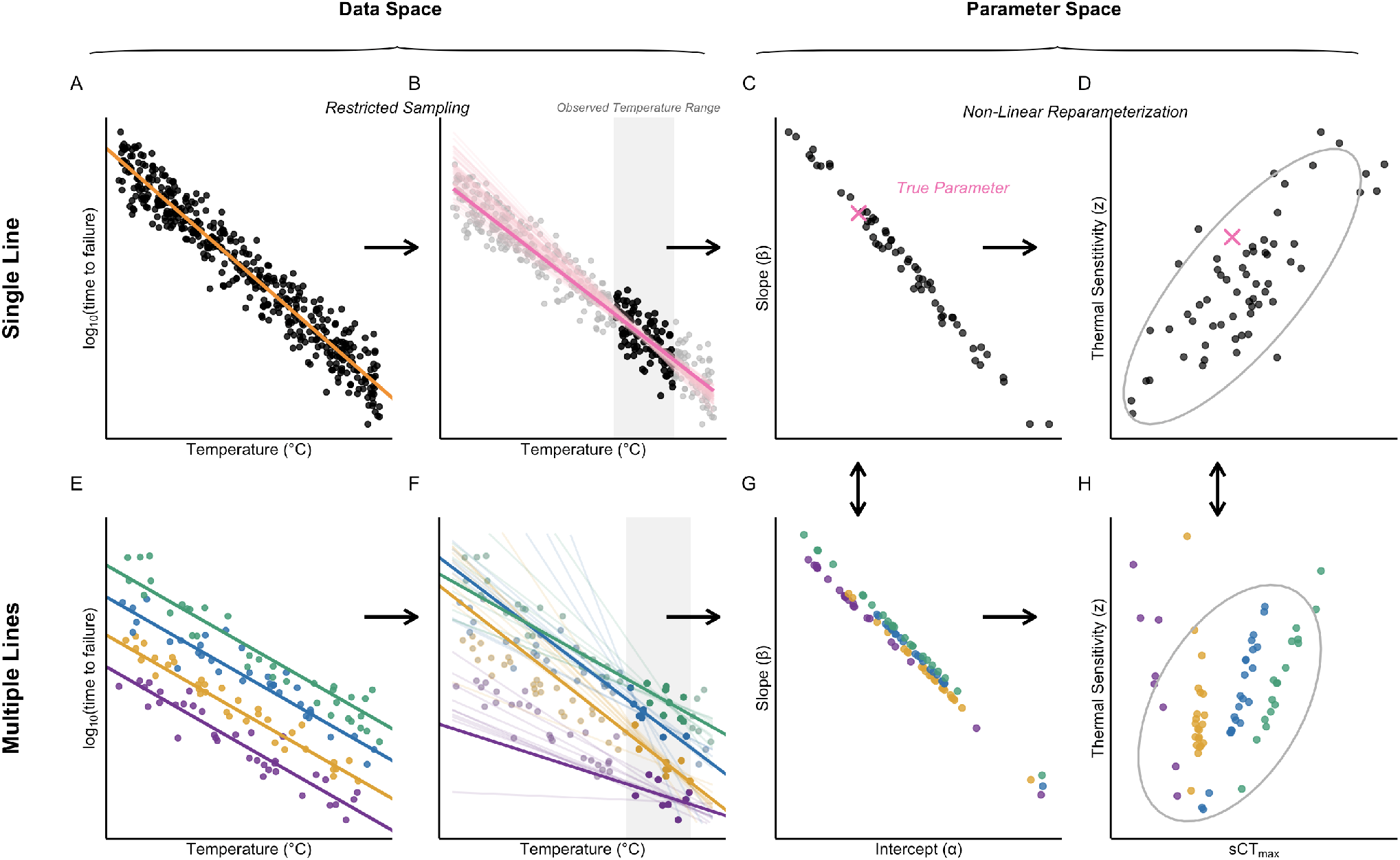
Mechanism of non-identifiability in thermal death time. **Panels A-C:** restricting sampling range results in a negative spurious correlation between estimated parameters (See Figure 1 for details). **Panel D:** Reparameterization to *z/sCT*_*max*_ results in a positive correlation. **Panels E-G:** The process from Panels A-C are repeated for multiple TDT lines (variation in parameters). **Panel H:** Reparameterization generates a positive correlation for each population, but these lines are not parallel, due to nonlinear reparameterization. See Figure S1 for demonstration with varying slopes.

### The artifact still emerges with trait variation

In practice, most traits vary at the individual, population, and species levels, and thermal tolerance is no exception (20, 26). We show that high induced correlation still emerges when considering several distinct TDT curves (representing thermal tolerance variation among multiple populations, species, etc.) (Figure 2E-H). However, the greater the difference between two generating curves, the muddier the *z/sCT*_*max*_ correlation between their generated data, as each TDT curve defines a distinct linear relationship between *z* and *sCT*_*max*_ (Figure 2E-H). Note that the TDT curves in Figure 2E are parallel for visual clarity, but the same trend emerges when slope is allowed to vary (Figure S1).

Quantifiable covariation in *α* and *β* (or *z/sCT*_*max*_) at a given biological level can be visualized with a joint probability distribution. In practice, we cannot know the “true” distribution, but we can estimate it by obtaining ordinary least-squares TDT fits for many individuals. However, Figure 2 showed that this estimation process produces compelling variation in TDT parameters when none truly exists–so how does this play out on a larger scale?

### Empirical TDT trade-offs can be generated by artifact

We have demonstrated the principle of non-identifiability in the general sense and within TDT. We now show that the population correlation structures observed in empirical TDT data can arise from this artifact alone. To do so, we compared empirical ordinary least-squares estimates of TDT curves from 100 isogenic lines of *Drosophila melanogaster* (Figure 3D, 3G) to estimates simulated under the null assumption of no biological signal (See methods) (Data: (21)). These Drosophila were sourced from the *Drosophila* Genetic Reference Panel (DGRP) (27).

**Fig. 3.**
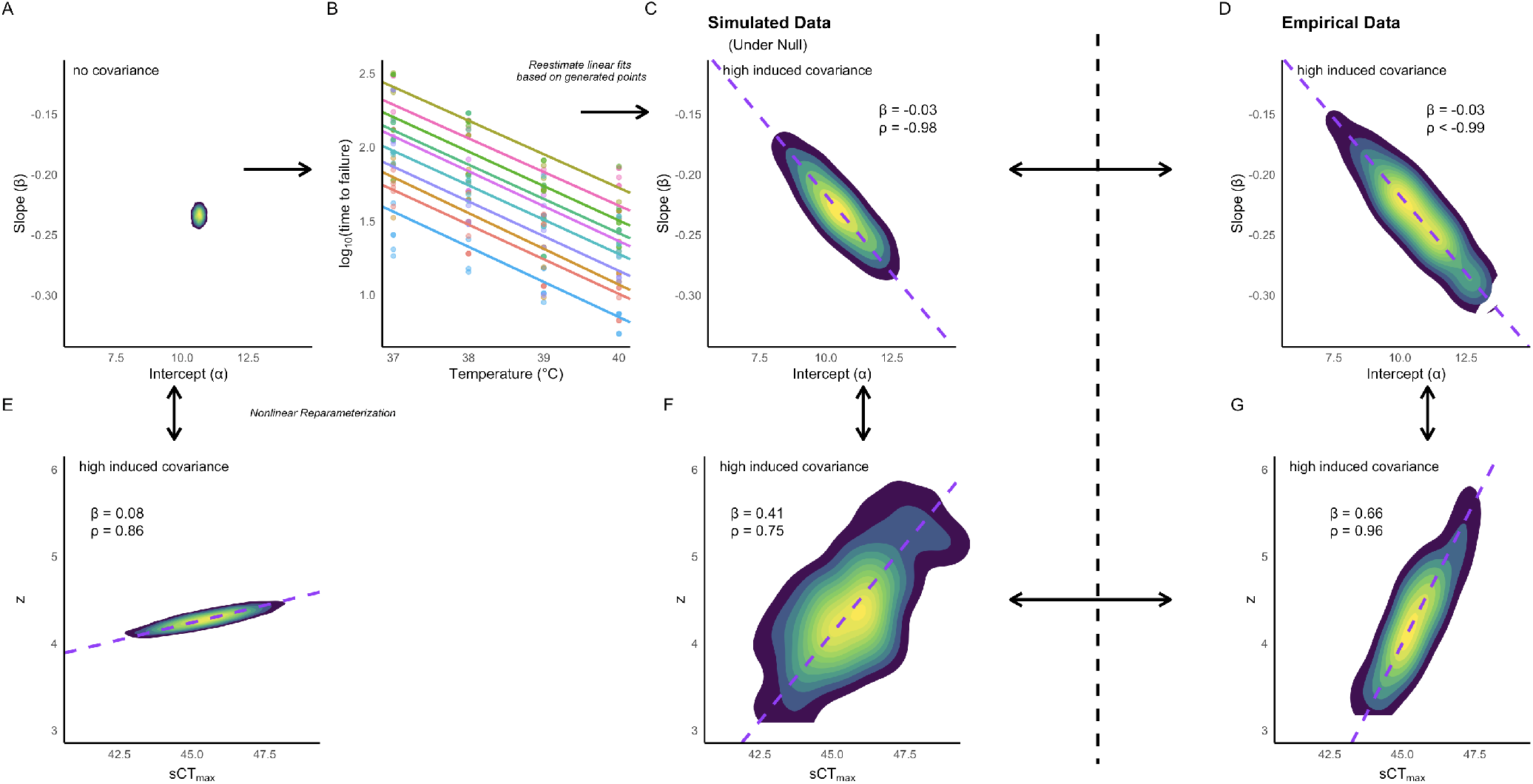
Empirical joint parameter structure mirrors artifact. **Panel A**: An uncorrelated generating joint distribution for slope and intercept, based on empirical estimates. **Panel B**: Slope/intercept pairs drawn from the distribution in panel A. Sampling mimicked protocol from the empirical dataset. 100 lines were simulated; we display 10 here. **Panel C**: Joint distribution of the 100 sampled lines. **Panel D**: Joint distribution of least-squares estimates for empirical TDT lines (Females). **Panels E-G**: Data from panels A, C, and D transformed into *z/sCT*_*max*_ space.

Comparing Figure 3C and 3D, we see that the joint distribution produced spuriously by sampling from an uncorrelated distribution is nearly identical to that of the empirical data (See Methods for details on the choice of null distribution). The slopes of their major axes are equal (−0.03), and their *α/β* correlations are nearly identical (−0.97 simulated vs < −0.99 empirical), although the normality assumptions imposed on the simulated data improves symmetry of the bivariate distribution (Figure 3C-D). With or without sampling error, reparameterizing the null distribution to *z/sCT*_*max*_ alters the correlation similarly to that observed in the literature (Figure 3E-G). Both *z* and *sCT*_*max*_ are functions of *−* 1*/β*. In fact, as *sCT*_*max*_ = *αz*, their correlation is a consequence of their premise and calculation. So, two processes: resampling error and non-linear reparameterization can both cause similar spurious correlations between *z* and *sCT*_*max*_. This simulation analysis shows that empirical correlations between traits can emerge from statistical artifact. We proceed with a more formal analysis to support this hypothesis.

### A framework for assessing trade-off identifiability

Building on the procedure demonstrated in Figure 3, we developed a methodology to determine the distribution of parameter correlations expected purely from non-identifiability defined by the data and model, with no biological signal. (details in Methods and SI). Using this technique with our *D. melanogaster* data showed that artifact alone could generate extreme correlations between slope and intercept (95% confidence interval: ♀: [−0.997, −0.993], ♂: [−0.995, −0.989]). Such extreme correlations could easily be mistaken for a biological trade-off under common practice, so we conclude that current evidence does not support the perceived TDT trade-off. However, this does not mean there is no trade-off; it simply means that it is not indicated by correlation alone. In fact, despite the magnitude of artifact, for both sexes, the observed correlation was significantly lower than expected by the null (♀: *r* = −0.999, *p* = 0.003;: −0.999, *p* = 0.001) (Figure S3). This result suggests that the correlation of the generating distribution may have been negative, supporting the trade-off hypothesis. Running the test for covariance instead of correlation yields similar results: (♀: 95% CI [−0.020, −0.012], cov = −0.050, *p* < 0.001; ♂: 95% CI [−0.016, −0.009], cov = −0.056, *p* < 0.001). We report the expected power under this testing and sampling procedure in Figure S4. Next, we use genomic data to confirm this finding that the TDT trade-off is real.

### Genomic signal in thermal tolerance parameters

Our above results show that estimated correlation and covariance are unreliable indicators of trait trade-offs. Here, we ran a Genome-Wide Association Study on the Principal component (PC) analysis of the *α/β* joint distribution for our *D. melanogaster* dataset to see if detectable genetic variation was present in the TDT trade-off axis (PC1) or net thermal tolerance axis (PC2) (Table 1).

**Table 1.** GWAS results for 4 parameter-phenotypes. Each cell contains the number of SNPs significant to the 10^*−*5^ level (DGRP standard–balancing multiple tests correction and realistic DGRP effect sizes (28)) for both the column and row parameters, for females/males, respectively. The diagonal contains the number of significant SNPs for a given parameter. 20 significant SNPs are expected by chance (~2,000,000 SNPs *∗*10^*−*5^). Dashes represent redundant cells.

| | PC1 | PC2 | $\beta$ | $\alpha$ |
| --- | --- | --- | --- | --- |
| PC1 | 79/28 | 0/0 | 77/25 | 72/26 |
| PC2 | — | 14/12 | 0/0 | 0/0 |
| $\beta$ | — | — | 98/27 | 71/24 |
| $\alpha$ | — | — | — | 75/27 |

For one or both sexes, PC1, *β*, and *α* are associated with a significantly greater count of SNPs than would be expected by chance (PC1: *p* < 0.001 ♀, *p* = 0.052 ♂*α*: *p* < 0.001 ♀, *p* = 0.078 ♂; *β*: *p* < 0.001 ♀, *p* = 0.078 ♂). Almost all SNPs significant to PC1 are significant to *α* or *β*, and vice versa, although slope was associated with slightly more SNPs than PC1 for females (Table 1). For males, PC1, *α*, and *β* are all associated with roughly the same number of SNPs (27–28), highlighting the highly unidentifiable nature of the TDT trade-off for the male dataset. This overlap was strong, but less severe for females, suggesting better identifiability. PC2 had no more significant SNPs than expected by chance (*p* = 0.934, ♀*p* = 0.979♂), and none overlapped with other parameters, showing that all detectable genetic variation was in the TDT trade-off (PC1) rather than net thermal tolerance (PC2). This result supports that the TDT trade-off is real, and carries serious implications for population adaptation under a warming climate.

We repeated this analysis under *sCT*_*max*_*/z* parameterization, returning a greater number of resulting significant SNPs in PC1 in comparison to *α/β* (♀/ ♂: 128/38), and again with a count of SNPs related to PC2 no greater than would be expected by chance alone (♀/ ♂: *p* = 0.895/0.052) (Table S1). We saw similar overlap between PC1 and its constituent parameters as in Table 1, although PC1 was related to more SNPs than *sCT*_*max*_ or *B* alone. The GWAS results for *B* and *z* were identical, as they contain the same mathematical information.

The purpose of this analysis was to draw inferences about biological signal captured by various parameterizations in unidentifiable systems–not to identify candidate SNPs associated with the thermal tolerance mechanism. The latter examination has already been done in detail for this dataset (21). Although some different candidate SNPs and alternative genetic analyses have arisen from our analysis, such details are beyond the scope of this study and remain avenues for future research.

### Non-identifiability in other modeling and biological systems

The non-identifiability phenomenon we have described is not rare–it is the norm when linear or non-linear models use limited data across a constrained range (11). Even within the context of grouped linear models, there are several possible modeling approaches, such as ordinary least-squares regression (e.g., TDT analysis above), linear mixed models (LMMs), and paired response estimation, that each come with their own sources of spurious correlation. We investigated datasets from the literature examining slope/intercept correlations or covariances in various biological applications to validate our test under these frameworks and to see how an artifact can influence results across biology.

### Reanalyzing trade-offs inferred from paired observation data

Sometimes the linear parameter estimates used to validate trade-offs are based on only two estimates: initial and subsequent. “Regression to the mean” (RTM), or the tendency for variability in initial response estimates to cause slope bias in the opposite direction, has been noted as an artifact with potential to produce spurious correlations and artificial trade-offs (8). Such a phenomenon can be considered a special case of non-identifiability, although no regression is involved. There exists a well-established method for correcting for RTM, and it has previously been applied to 25 datasets that reported trade-offs in initial thermal tolerance (either cold or heat) and plasticity (a distinct trade-off from TDT), finding that only 6 remained significant after correction (16, 29–40). Comparison to this meta-analysis provides an ideal way to validate our test.

We ran the test (described above) on these 25 datasets (details and code in SI). In every instance, the null correlation was negative (Figure S6, Table S2). The observed empirical correlation was significantly more negative than expected under the null for 4/25 datasets (all heat tolerance), while it was greater (less negative, or more positive) for 2 (both cold tolerance). The two techniques (RTM corrected correlation, and trade-off test) agreed on result significance in 20/25 cases. In 22/25 cases, the sign of the adjusted correlation matched the relationship of the empirical correlation to the median null correlation (e.g., positive correlation, empirical greater than null). 2/3 of the non-matching cases had |*r*| > 0.05. Qualitatively, when adjusted correlation was near zero, the observed correlation was near the median under the null (Figure S6).

### Reanalysis of linear mixed model trade-offs

To determine how non-identifiability affects correlation estimates differently in a linear mixed modeling framework, we used our framework to analyze 3 linear mixed model reaction norm studies that interpreted a slope-intercept correlation as a trade-off.

### Passerine bird aggression trade-off

Previous researchers analyzed the aggressiveness of mature male Great Tits (*Parus major*) across breeding stages (breeding and non-breeding), controlling for random effects of individual and brood (Figure S7A) (41). They found a correlation between the resulting per-individual slopes and intercepts (even after centering) (Figure S7B), interpreting a possible trade-off between overall aggressiveness and plasticity of aggressiveness. Here, we apply our identifiability test, finding that the null distribution of parameter correlations expected under artifact is apparently unbiased, with the null being centered around 0 (95% confidence interval: (−0.24, 0.25). The observed trade-off is statistically significantly lower than would be expected under the null (*r* = −0.681, *p* = 0.002).

### Stickleback behavioral trade-off

It has been claimed that the plasticity (slope over time) of boldness, or “tendency to explore an open arena from the safety of cover,” of the threespined stickleback (*Gasterosteus aculeatus*) is in a trade-off with their boldness at the beginning of their time in that arena (intercept) (*r* = −0.56) (Figure S7D-E) (42). Our simulation shows that the correlation of these two variables is poorly constrained with a strong right skew under the LMM and data structure (95% confidence interval: (−0.4, 0.92) (Figure S7F). We did not center the predictor for this test, adhering to the original analysis. Despite the unconstrained null, the observed correlation was below what would be expected under artifact alone (*r* = −0.56, *p* = 0.002).

### Mussel sperm motility vs longevity reanalysis

Sperm motility has been proposed to hold a trade-off with its longevity, indicated by a negative correlation between the two traits. This was tested in the broadcast-spawning mussel, *Mytilus galloprovincialis*, where the researchers interestingly found the opposite effect: a positive correlation between the LMM intercept (baseline motility) and slope (longevity) (*r* = 0.79, *p* = 0.064). Our simulation shows that the correlation of these two variables is poorly constrained (95% CI: (−1,1)) (see Discussion for explanation), but unbiased (Figure S7I). Thus, we fail to reject the null hypothesis that the observed correlation arose from an uncorrelated generating distribution (*p* = 0.264).

### Analytical expressions show ways to mitigate non-identifiability

We derived analytical expressions for the effects of sampling design and residual error on estimated parameter variance and covariance (Table S3; SI Text). These expressions show that centering eliminates “structural,” or model-driven, slope-intercept covariance. Centering our *Drosophila* data demonstrates this finding empirically, as the centered *cor*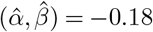for females and −0.41 for males: weaker than the <-0.99 estimated from non-centered data (S2). After centering, the remaining non-identifiability is “practical,” or inherent to the dataset, and it decreases with greater replication and predictor variance. Simulations confirmed these predictions and showed that, for a fixed sample size, two widely separated predictor values improve identifiability over many intermediate values (Figure S8).

## Discussion

### Function-valued trait trade-offs under the lens of non-identifiability

When biological traits are represented by statistical parameters, their estimated properties include two distinct sources of influence: genuine biological signal, and statistical structure imposed by the data structure and model design. When several combinations of model parameters describe variation in a model similarly well, parameter estimates form a predictable joint pattern in parameter space independently of underlying signal in the data–when unrecognized, this passive structure can be harmful, as it gives the illusion of biological signal when none exists, resulting in potentially misleading conclusions across the literature. Such an artifact can arise from many sources: “ridge” geometry arising from poorly identified linear models (Figure 1), “shrinkage” imposed by mixed linear modeling software (Figure S7; more below), nonlinear reparameterizations, relating ratios of the same variable (Figure 3) (2), failing to center data (Figure S2, Figure S7D-F), regression to the mean (Figure S6) (1, 8, 16), and more. While past trait biology literature has tended to treat each of these problems as independent, here, we show that it is useful to consider them all under the broad lens of non-identifiability. Artifactual sources of parameter signal make it ill-advised to interpret statistical parameters as traits without first understanding the influence of model and data structure on their estimation. While this is especially true for the correlation and covariance parameters we focused on here, structural bias in other parameters such as variance can be strong (Figure 1, Figure 3, Table S3).

We demonstrate these points by exploring the correlation between Thermal Death Time sensitivity and upper critical threshold, which is commonly interpreted as a trade-off, showing that lack of centering, nonlinear parameterization, relating two variables with a common denominator, and linear ridge geometry all play roles in biasing estimations of this parameter (Figure 2, Figure 3, Figure S1, Figure S2). In fact, we use simulation from an uncorrelated null distribution to show that correlations as high as −0.99 could be expected in the absence of true biological covariance (Figure 3, Figure S3, Table S2). However, we repeat this process many times to form a null distribution for correlations expected under artifact alone, and compare the observed correlation to this distribution in a statistical test, showing that in fact, biological covariance was likely necessary to achieve the observed correlation (Figure S3). This finding was supported by the results of a GWAS, showing that genetic variation was present in the trade-off axis (PC1) of the TDT slope-intercept joint distribution, showing that the trade-off is in fact biological (Table 1) (See Discussion for axis interpretation). At the same time, we found that no genetic variation was present orthogonal to this axis, providing evidence that net thermal tolerance is highly conserved within this population of *D. melanogaster*, and thus it has little to no potential to adapt to a warming climate (Table 1). In addition to strong thermal tolerance and conservation implications, these results show that trait trade-offs inferred from correlation data can be real, and that statistical methods can be used to extract this information from even extremely non-identifiable parameters.

Biological interpretation of these trade-offs is context-dependent, and even after validating a statistical correlation in the true biological distribution of traits, researchers must consider biological phenomena other than true trade-offs that may cause such correlations (i.e., “ghost trade-offs”), and the complex resource allocation shape and structure that may govern the strength of true trade-offs (43, 44).

### Generalizing the framework

Having developed and demonstrated methods for extracting biological information from poorly identifiable data in the context of TDT, we broadened our scope to explore the implications of non-identifiability in other biological and statistical contexts. Our reanalysis of 25 paired observation trade-off datasets showed that our proposed framework agrees with established artifact correction techniques in a specific case where such a method exists (Table S2). In cases where the corrected correlation was around 0, our observed correlation was around the null median; when the corrected correlation was negative, we observed a correlation less than the null median. However, our procedure tended to estimate slightly less negative (or more positive) biological correlation than the ROM correction, possibly due to correlation bias such as centering bias that our general method accounts for, but the established method missed.

Analysis of trade-off datasets modeled using LMMs showed that the negative bias present in OLS regression is not present using the R function *lmer* for hierarchical modeling. Rather, these unbiased correlation and covariance distributions (Figure S7, C, F, I) are products of more complicated sources of artifact that constrain the median to 0, push poorly identifiable fits to both positive and negative extremes near 1 and −1 (e.g., Figure S7F, I), and simultaneously shrink covariance estimates towards 0, although these trends may change with model structure (45, 46). Thus, while the mean of this distribution is unbiased, its variance is not, leaving LMM correlations especially difficult to interpret biologically. However, our simulation procedure does not interpret the magnitude of these biased estimates, nor does it require explicit inclusion of each complex artifact. Rather, it interprets the position of the observed result relative to these estimates. So, a significant result from our test still yields the same conclusion under the context of LMMs: the observed correlation was more extreme than expected under the null of no covariance. As such, the framework is not model-specific.

For more complex, even non-linear models, the procedure for testing the presence of biological signal in your parameter estimates can remain the same: 1) Define a null parameter constraint (here: slope/intercept covariance=0). This places the assumption that the true biological signal in the data follows this constraint. 2) Estimate the model under this constraint. This removes influence from the model/data structure on this parameter, giving a reasonable estimate for a biological parameter distribution under the null. 3) Sample from the distribution defined by this model, following your empirical sampling procedure (same predictors). Refit your model (without the constraint) on this model, and obtain your parameter estimate. This emulates empirical procedure: collect your data (with variability) and fit your model to that data. This step imposes data/structure bias on your parameter estimates. 4) Repeat (3) many times, forming a null parameter distribution. 5) Compare your observed parameter to this distribution. Note that the parameter constraint is ambiguous: variance, mean, or other constraints may be relevant depending on biological context and expected artifacts within your modeling procedure.

### Suggested analytical and empirical practices

Model identifiability can be addressed in three ways: diagnosis, analytical mitigation, and experimental design. A routine check for flagging potentially poor model identifiability is checking for correlation/covariance between the distribution of parameter estimates. Parameter variance/covariance matrices are produced by many statistical modeling tools, such as *lm/glm* in *R*. If poor identifiability is expected, there are many ways to diagnose it. Our framework test offers a good way to assess the constraint and identifiability of a model and its parameters, indicated by the variability in the null distribution.

Analytical techniques can rescue “structural” non-identifiability, which is coordinate-system dependent. Notably, centering data and good variable choice can reduce, or in the case of simple linear models, eliminate this bias (Table S3; Figure S2, Figure S7). However, after centering, some artifact often remains (e.g., ridge geometry outlined in Table S3). This “practical” non-identifiability is inherent to the dataset, but it can be reduced *a priori* through conscious experimental design. Increasing sample size, maximizing predictor variance, and minimizing sampling error can all improve the identifiability of a dataset. While we mainly explored correlation in this work, we encourage the use of covariance over correlation for trade-off analysis in many scenarios, as it contains more information regarding the scale of the parameter distributions and is not constrained to the range (−1,1), making its distribution more interpretable (e.g., Figure S3, Figure S6, Figure S7). Our test functions in *R* are interchangeable between correlation and covariance (47).

It is necessary to assess the sample size and predictor range needed to improve identifiability to desirable levels. This type of power analysis can greatly increase data quality relative to resource expenditure (if done *a priori*), and can put results in statistical context (if done *a posteriori*). We performed such an analysis based on our *Drosophila* TDT data, showing that for this empirical sampling structure, nearly 1000 reaction norms (curves relating the phenotype of a sample population to an environmental variable–e.g., a TDT curve) are required to generate a power of >0.8, given a moderate true slope-intercept correlation of *r* = 0.5/-0.5 (Figure S4). These estimates are approximations, as the parameters of the true generating distribution are unknowable (but estimable); however, they can provide a reasonably conservative baseline.

While 1000 reaction norms are an extreme and often impractical sample size for most situations, our results emphasize the importance of sampling design in dictating the power associated with a reaction norm study: this TDT dataset has equally spaced predictors across a narrow range (3°C). Following our above sampling design suggestions could reduce the necessary sample size considerably. Our results emphasize how statistical power can vary greatly for a given sample size, agreeing with previous analyses (48, 49). However, we provide important nuance to the previously suggested rules of thumb of n=200-1000 samples (individuals, not reaction-norm curves) in reaction norm analysis, which did not account for non-identifiability or emphasize the importance of underlying parameter structure, and thus likely overestimated the statistical power and underestimated the necessary sample size of many reaction norm analyses, (48, 49). As power varies greatly based on statistical context, we cannot support a single “rule of thumb” sample size, and rather encourage thorough context-specific power analysis. We provide a reproducible function, pairing such a power analysis with our previously described test, for researchers to use for their own purposes (See Methods, Supporting Code). Analytical expressions (e.g., Table S3) or simulation analyses (e.g., Figure S8) are alternative avenues, and online tools also exist for this purpose (see (50)). Researchers should account for the fact that failure to propagate uncertainty can bias reaction norm power and interpretation (51). While this is a separate problem, it could be more consequential for poorly identifiable parameters with large uncertainty.

Finally, to produce interpretable parameters, model parameterization should reflect the effective dimensionality of the data. Reparameterization or dimension reduction may improve interpretation (e.g., Table 1), whereas parameters with shared components, such as *z* and *sCT*_*max*_, can induce additional correlation (Figure 3) (2). Poorly identified parameters should therefore not be interpreted as independent traits without supporting evidence. For example, PC1 of our uncentered TDT slope/intercept joint distribution explained >99% of variation in these parameters for both males and females, indicating that *α* and *β* effectively jointly describe a single trait (the trade-off) rather than two independent traits (Figure S5).

Our *Drosophila* dataset contained 100 independently fitted least-squares curves, allowing us to estimate mean parameters well while revealing that variance and covariance were dominated by estimation error. However, many studies lack such replication and may only estimate one model. Such estimates could lie anywhere on the non-identifiability ridge, producing large parameter uncertainties without the replication to diagnose the issue. In such cases, it is especially important that researchers take necessary precautions to identify and mitigate poor identifiability *a priori*.

Trade-offs are already notoriously difficult to detect due to the heterogeneity in resource distributions and allocation pathway structures (43), and consideration of spurious correlations shows that they are likely more elusive than previously realized. However, the tools we outline here provide ways for researchers to more easily extract trade-off signal from statistical noise, avoid erroneous interpretation of an artifactual correlation, plan sampling structure to maximize resources, and assess the level of confidence they should have in a null result.

### Use of Parameters as Traits

Our results support caution against the use of parameters as traits without assessing their pitfalls and limitations (2–4, 52). It has been argued that functions as a whole are themselves traits under selection, while parametrically-derived parameters are often simply researcher-defined components for describing these complex phenomena and may not correspond to biological traits themselves (53, 54). Through the lens of identifiability, such a conclusion seems reasonable, as poorly identified models can still yield highly constrained and valuable results, even if their parameters are uninterpretable (11). For instance, TDT models sharing a non-identifiability “ridge” describe the same net thermal tolerance outcome, despite widely variable parameter estimates, giving our TDT data an effective dimensionality of one, although the model is typically described by two parameters (Figure 2 and Figure 3). However, we also show a very clear dichotomy between the genomic signal in PC1 and PC2 (Table 1), indicating that these parameters are very much biologically independent. Similarly, we were able to detect clear biological signal in correlation and covariance, independently of function-wide effects on these parameters, and corroborate that signal with the aforementioned genomic data. Thus, our results support that some parametric parameters can indeed be considered independent biological traits, only if appropriate steps are taken to sift that signal from model-wide noise. Specifically, we show that a trade-off itself can be defined by a parametrically derived parameter, driven by a constraint on the function-valued response trait.

### Implications for thermal tolerance

The extreme correlation between thermal sensitivity (*z* =*−* 1*/β*) and critical thermal limit (*sCT*_*max*_ =*− α/β*) can be expected under statistical artifact alone (Figure S3). Specifically, correlated estimation error in *α* and *β* translates through the non-linear reparameterization to *z* and *sCT*_*max*_, whose shared denominator introduces additional correlation, while lack of centering accentuates this issue (Figure 2, Figure 3, Figure S2, Figure S3). However, our simulation test shows that the observed TDT correlation (in *α/β* space) is stronger than expected under artifact alone, suggesting that the trade-off is real. Despite this, observed variation and covariation in *sCT*_*max*_ and *z* (or *α/β*) is likely greater than its biological magnitude (Figure 2A vs 2D). Next, we justify the genetic analysis that further supports our trade-off claim.

Any biological trade-off in parameter space would likely occur along the same axis as the artifact, as both statistical and biological trade-offs are defined by the spectrum of parameter combinations that achieve a similar response within the sampled range. We therefore interpret PC1 as this “trade-off axis,” while PC2 captures variation in net thermal tolerance: towards or away from the ideal “jack of all trades” phenotype (Figure 3). TDT lines with relatively higher average death times are displaced in a common direction of PC2 (top-right) (Figure 2E-G, Figure S2), while their shared data structure defines parallel uncertainty ridges in PC1. However, this trend is obscured in *z/sCT*_*max*_ parameterization (Figure 2H), making *α/β* the preferred coordinate system for separating trade-off strategies from net thermal tolerance.

Our GWAS results show significant biological signal along the trade-off ridge (PC1), but none in net thermal tolerance (PC2), showing that all detectable genetic variation was among trade-off strategies, not absolute thermal tolerance within the sampled range (Table 1, Table S1). The substantial overlap in significant SNPs described by *α, β*, and PC1 (as well as *sCT*_*max*_*/z*) further indicates that typical TDT parameters describe a shared biological axis system rather than independent traits.

These findings support the trade-off hypothesis for thermal tolerance and suggest that the magnitude of thermal tolerance determined by genetics is highly conserved for these lines of *D. melanogaster*. These results are striking in the face of climate change: we show that detectable genetic variation in this population of *Drosophila* is limited to this trade-off axis rather than an unambiguous increase in overall thermal tolerance. In other words, this population may lack the genetic variation to adapt to a warming climate. While these standardized environment experiments do not account for gene-by-environment interactions or plasticity, they could be highly informative for projecting biotic responses to climate change if replicated for other taxa, and our pipeline shown here provides a way to do so. Past studies have found genetic variation in thermal tolerance, and even evidence of adaptation under a warming climate (55, 56). However, many of these studies use linear ramping *CT*_*max*_ (not *sCT*_*max*_) as a thermal tolerance metric: a 1-dimensional trait that cannot distinguish between trade-off and net thermal tolerance (57). Other studies demonstrate that TDT parameters show genetic variation, a result that is consistent with PC1 being defined by these parameters (20, 58). These studies show that selection on one TDT parameter induces changes in the other TDT parameter (likely via PC1), further supporting our conclusions. More broadly, studies have repeatedly found genetic variation in reaction norm plasticity (slope) and initial value (intercept) (59), but our results show that PC decomposition is necessary to determine whether this variation was directly in the reported traits, or in a trade-off between the two. Our results support and provide nuance to previous claims that plasticity, not genetics, shapes individual responses to thermal stress (60). Analysis of TDT PC variation with respect to acclimation temperature is an area for future work. The fitness implications of high vs low PC1 values, and the reason for the highly conserved nature of net thermal tolerance remain to be seen. Models that predict ectotherm survival under fluctuating temperature conditions may offer good tools for assessing the selection gradient of this phenotype across variable climates (e.g., (19, 57, 61–64)).

### Preferred TDT parameterization

TDT models can be parameterized by least-squares intercept (*α*) and slope (*β*) or by defined parameters *z* and *sCT*_*max*_. In theory, *sCT*_*max*_ provides a more interpretable parameter than *α* for comparison across populations. *sCT*_*max*_ (or *sCT*_*min*_) may be more identifiable than *α* when *sCT*_*max*_ is closer to the observed temperature range, and we saw many more significant SNPs associated with *sCT*_*max*_ than *α*, which could have been a product of better identifiability, non-linearity, or increased biological signal in *sCT*_*max*_ (Table 1, Table S1). However, centering makes *α* much more interpretable (average *log*(*t*_*f*_) at a central temperature) and eliminates structural identifiability problems. While *sCT*_*max/min*_*/z* parameterization defines a line, there is no way to directly linearly estimate either parameter while preserving the error structure of the TDT model. These nonlinear transformations change the data from its natural linear coordinate system, making interpretation less intuitive and statistical analysis (e.g., PCA) more complicated (see Figure 2, Figure S1). Additionally, *z* and *sCT*_*max*_ are indirectly estimated and explicitly linked (*sCT*_*max*_ = *zα*), which causes inherent correlation, regardless of biological signal or dataset identifiability (Figure 3). Although *β* and *z* contain the same mathematical information and the same genetic information in our study (Table S1), transformation to *z* can distort analyses that depend on scale or linearity. Thus, we recommend *α/β* parameterization for analyzing or interpreting parameter distributions, while *sCT*_*max/min*_ is reserved as a summary statistic when its biological interpretation is useful, especially for inter-dataset comparison where *α* may be centered at different temperatures. Note that use of *sCT*_*max/min*_ should be paired with response (*log*(*t*_*f*_)) centering rather than predictor (temperature) centering.

We advocate for biologists to make identifiability checks and data centering part of their modeling routine, and we encourage those designing experiments to consider *a priori* how their sample size, predictor range, and other aspects of study design will influence the identifiability and statistical power of their analyses. In this work, we considered how model identifiability and statistical artifacts affect biological inference through the lens of TDT. In doing so, we made several inferences with broad implications for thermal biology under a warming climate. We hope that this work inspires and enables researchers in many subfields to do the same, improving the quality of parameter inference across biology. Broadly, we emphasize that model parameters should not be interpreted as independent biological traits, nor should correlations among them be taken as evidence of biological trade-offs, until the information supporting them and the statistical biases influencing them have been thoroughly examined.

## Materials and Methods

### Null model selection

To demonstrate that the joint distribution of TDT parameters could qualitatively resemble one generated by artifact, we simulated population TDT data from under the null assumption that *cor*(*α, β*) = 0 for a hypothetical population. We defined a known null population distribution of parameters *α, β*, and error *ϵ*, where we defined *cor*(*α, β*) = 0, and we mirrored the dataset’s empirical protocol by drawing 100 pairs of *α* and *β* from this null distribution (100 isogenic lines with varying slope and intercept), similarly to (2). We then generated observations with randomly sampled error from each of these lines at 4 temperatures (37-40°C) (assuming a shared error distribution across lines) and reestimated the least-squares fit of each of these 100 groups, obtaining an estimated joint distribution of *α* and *β* that mirrored the empirical data.

We chose the null distribution through an optimization process to produce the estimated joint distribution of *α* and *β* closest to the empirical data. We repeated the above simulation using many uncorrelated null distributions, allowing the variances of *α, β*, and *ϵ* to vary across a grid, while their means were fixed to empirically estimated values (male and female data were combined for this qualitative assessment). For each candidate null, we compared the resulting estimated joint *α/β* distribution to the empirical distribution using a sum of squared errors metric. The null distribution that minimized this metric was selected for visualization.

### Artifact test

We repeated the above sampling procedure many times, creating a null distribution of correlations or co-variances expected under artifact alone. We made this procedure accessible in a generalized and reproducible function (see Supporting code) that modifies the optimization and resampling procedure based on provided model and data structure. Here, the optimization structure was more formal than in Figure 3, resulting in slightly different correlation estimates (see SI for details). A second such function applies the same procedure to mixed effects models defined by the *lmer* function in *R* (*lme4* package) (65). *Lmer* allows for explicitly defined null models with parameter covariance of 0, so we use the optimization procedure in this package, with the same resampling and reestimation steps as the ordinary least squares/paired observation example. We compare the empirically observed correlation or covariance to the generated null distribution to estimate a p-value. Our null hypothesis is that all observed correlation or covariance is due to model and data structure, and none is biological. As such, we use a two-tailed test to detect positive and negative deviations. If *p* <0.05, we can conclude that something other than artifact (e.g., biological effect) is influencing the observed correlation or variance/covariance structure. While our study only uses this test to detect spurious correlations or covariance, an analogous procedure could test for signal in parameter variance, or other moments.

We developed a reproducible function in *R* to assess the power of our test under a given sampling protocol and a given generating distribution with a defined correlation (Supporting code). This procedure involves sampling (with error) from a defined generating distribution (i.e., the “true” biological distribution) according to a sampling procedure defined by the user. Then, the trade-off test is applied to this sampled data, and we record whether a significant result is obtained. We repeat this procedure many times to obtain a statistical power/sensitivity for a range of possible correlations. While this function is designed for ordinary least-squares compatibility, the same procedure could be followed for use with *lmer* models. We demonstrated this power analysis in the context of the isogenic *Drosophila* TDT data, estimating power as a function of biological correlation and reaction norm sample size (Figure S4).

### Dataset and GWAS analysis

The dataset used in this study was obtained from (21). See that study for details regarding fly rearing, TDT data collection, and genome sequencing. The dataset consisted of 100 isogenic lines of *Drosophila melanogaster* from the *Drosophila* Genome Reference Panel (27), with 5 flies sampled per line per temperature (37, 38, 39, 40°C) per sex. Slopes (*β*) and intercepts (*α*) were obtained via ordinary least-squares regression for each isogenic line and sex. PC1 and PC2 were calculated in *R* for each line and sex by running PCA on the joint distribution of *β* and *α* (code in SI). Phenotypes *α, β, z, sCT*_*max*_, PC1 and PC2 were uploaded to DGRPool, an online tool for standardized DGRP GWAS analysis (66). See that work for data filtering and analysis standards. We ran a GWAS on these phenotypes, counting the number of SNPs, for each combination of phenotype, line and sex, with p-values below the DGRP standard significance threshold of 10*−*5, which balances a correction for multiple tests with realistic levels of signal typical of significant DGRP SNPs (28).

### Linear Mixed-Model Dataset Selection

We selected three datasets on the following conditions in the original publications: 1) they interpreted a slope-intercept correlation as a biological trade-off 2) they used LMM approaches using *lmer* in *R*, and 3) they made their full dataset and modeling procedure/code publicly available. While the search was not exhaustive or fully systematic, no other datasets were analyzed, and no datasets were excluded after selection.

### AI use

Generative AI (Microsoft 365 Copilot, enterprise version) was used for assistance with writing (editing, formatting), some coding (debugging, formatting, generation), and mathematical validation. The authors reviewed, validated, and synthesized output, and are responsible for all content.

***See Supporting Information for more detailed methodology not addressed here, including supporting figures and text***.

## Code and data availability

All code and data associated with this work can be found at https://github.com/gbullard87/Tradeoff_artifacts_2026

## Acknowledgments

We thank Mete Yuksel for advice regarding genomic analyses, Nicole Bison for discussions related to TDT parameter correlation, and Erik Curtis for comments on previous versions of the manuscript.

## Bibliography

1. Nils Blomqvist. On the bias caused by regression toward the mean in studying the relation between change and initial value. Journal of Clinical Periodontology, 14(1):34–37, 1987. doi: 10.1111/j.1600-051X.1987.tb01510.x.

2. D. A. Jackson and K. M. Somers. The spectre of “spurious” correlations. Oecologia, 86(1): 147–151, 1991. doi: 10.1007/BF00317404.

3. T. Ergon and R. Ergon. When three traits make a line: evolution of phenotypic plasticity and genetic assimilation through linear reaction norms in stochastic environments. Journal of Evolutionary Biology, 30(3):486–500, 03 2017. ISSN 1010-061X. doi: 10.1111/jeb.13003.

4. Jip J. C. Ramakers, Marcel E. Visser, and Phillip Gienapp. Quantifying individual variation in reaction norms: Mind the residual. Journal of Evolutionary Biology, 33:352–366, 2020. doi: 10.1111/jeb.13571.

5. Kenneth A. Bollen. Structural Equations with Latent Variables. Wiley, 1989.

6. Douglas S. Falconer and Trudy F. C. Mackay. Introduction to Quantitative Genetics. Longman, 4 edition, 1996.

7. Roosa A. E. Laitinen and Zoran Nikoloski. Strategies to identify and dissect trade-offs in plants. Molecular Ecology, 33(10):e16780, 2024. doi: 10.1111/mec.16780.

8. Colleen Kelly and Trevor D. Price. Correcting for regression to the mean in behavior and ecology. The American Naturalist, 166(6):700–707, 2005. doi: 10.1086/497402.

9. Enrico L. Rezende and Francisco Bozinovic. Thermal performance across levels of biological organization. Philosophical Transactions of the Royal Society B: Biological Sciences, 374(1778):20180549, 06 2019. ISSN 0962-8436. doi: 10.1098/rstb.2018.0549.

10. Jacob A. Moorad. Mathematical artifacts in slope–intercept correlations invalidate biological inferences in aging research. Proceedings of the National Academy of Sciences, 122(50):e2523907122, 2025. doi: 10.1073/pnas.2523907122.

11. Ryan N. Gutenkunst, Joshua J. Waterfall, Fergal P. Casey, Kevin S. Brown, Christopher R. Myers, and James P. Sethna. Universally sloppy parameter sensitivities in systems biology models. PLoS Computational Biology, 3(10):e189, 2007. doi: 10.1371/journal.pcbi.0030189.

12. Tjalling C. Koopmans and Olav Reiersøl. The identification of structural characteristics. The Annals of Mathematical Statistics, 21(2):165–181, 1950. doi: 10.1214/aoms/1177729394.

13. Linda Wanika, Joseph R. Egan, Nivedhitha Swaminathan, Carlos A. Duran-Villalobos, Juergen Branke, Stephen Goldrick, and Mike Chappell. Structural and practical identifiability analysis in bioengineering: a beginner’s guide. Journal of Biological Engineering, 18:20, 2024. doi: 10.1186/s13036-024-00410-x.

14. Charity J. Morgan. Use of proper statistical techniques for research studies with small samples. American Journal of Physiology, 313:L873–L877, 2017. doi: 10.1152/ajplung.00238.2017.

15. Alexander P. Browning, Christopher Drovandi, Ian W. Turner, Adrianne L. Jenner, and Matthew J. Simpson. Efficient inference and identifiability analysis for differential equation models with random parameters. PLoS Computational Biology, 18(11):e1010734, 2022. doi: 10.1371/journal.pcbi.1010734.

16. Alex R. Gunderson. Trade-offs between baseline thermal tolerance and thermal tolerance plasticity are much less common than it appears. Global Change Biology, 29(13):3519– 3524, 2023. doi: 10.1111/gcb.16710.

17. Enrico L. Rezende, Luis E. Castañeda, and Mauro Santos. Tolerance landscapes in thermal ecology. Functional Ecology, 28(4):799–809, 2014. doi: 10.1111/1365-2435.12268.

18. Luis E. Castañeda, Enrico L. Rezende, and Mauro Santos. Heat tolerance in drosophila subobscura along a latitudinal gradient: Contrasting patterns between plastic and genetic responses. Evolution, 69(10):2721–2734, 2015. doi: 10.1111/evo.12757.

19. Enrico L. Rezende, Francisco Bozinovic, András Szilágyi, and Mauro Santos. Predicting temperature mortality and selection in natural drosophila populations. Science, 369(6508):1242–1245, 2020. doi: 10.1126/science.aba9287. PMID: 32883867.

20. Andrés N Molina, Mauricio J Carter, and Enrico L Rezende. Plasticity cannot fully compensate evolutionary differences in heat tolerance across fish species. Evolution, 78(12):1949–1957, 12 2024. ISSN 0014-3820. doi: 10.1093/evolut/qpae126.

21. J. Soto, F. Pinilla, P. Olguín, and L.E. Castañeda. Genetic architecture of the thermal tolerance landscape in drosophila melanogaster. Molecular Ecology, 34(7):e17697, 2025. doi: 10.1111/mec.17697. Epub 2025 Mar 4; PMID: 40035350.

22. Larry Wasserman. All of Statistics: A Concise Course in Statistical Inference. Springer, New York, 2004. ISBN 9780387402727.

23. Andreas Raue, Clemens Kreutz, Thomas Maiwald, Julie Bachmann, Marcel Schilling, Ursula Klingmüller, and Jens Timmer. Structural and practical identifiability analysis of partially observed dynamical models by exploiting the profile likelihood. Bioinformatics, 25(15):1923–1929, 2009. doi: 10.1093/bioinformatics/btp358.

24. Helena S. Bayat, Sebastian Prati, and Ralf B. Schäfer. Synthesizing evidence to beat the heat: a guideline for thermal tolerance reporting. Journal of Thermal Biology, 136:104432, 2026. ISSN 0306-4565. doi: 10.1016/j.jtherbio.2026.104432.

25. W. D. Bigelow. The logarithmic nature of thermal death time curves. The Journal of Infectious Diseases, 29(5):528–536, 1921. doi: 10.1093/infdis/29.5.528.

26. Scott Bennett, Carlos M. Duarte, Núria Marbà, and Thomas Wernberg. Integrating within-species variation in thermal physiology into climate change ecology. Philosophical Transactions of the Royal Society B, 374(1778):20180550, 2019. doi: 10.1098/rstb.2018.0550.

27. Trudy F. C. Mackay, Stephen Richards, Elissa A. Stone, Antonio Barbadilla, Julien F. Ayroles, Dongsheng Zhu, Sonia Casillas, Youngseok Han, Michael M. Magwire, Julie M. Cridland, Matthew F. Richardson, Robert R. H. Anholt, Manel Barrón, Celeste Bess, Kerstin P. Blankenburg, Matthew A. Carbone, David Castellano, Laurie Chaboub, Laura Duncan, Zachary Harris, Muhammad Javaid, Jothi C. Jayaseelan, Shalini N. Jhangiani, Kurt W. Jordan, Federica Lara, Fred Lawrence, Song Yee Lee, Pablo Librado, Raquel S. Linheiro, Richard F. Lyman, Andrew J. Mackey, Manraya Munidasa, Donna M. Muzny, Leena V. Nazareth, Ifunanya Newsham, Laura Perales, Li L. Pu, Chunhui Qu, Miriam Ramia, Jamie G. Reid, Stephanie M. Rollmann, Julio Rozas, Naïr Saada, Laxmi Turlapati, Kevin C. Worley, Yang Q. Wu, Akihiko Yamamoto, Yufeng Zhu, Casey M. Bergman, Kevin R. Thornton, David Mittelman, and Richard A. Gibbs. The drosophila melanogaster genetic reference panel. Nature, 482(7384):173–178, 2012. doi: 10.1038/nature10811.

28. Paul Battlay, Pontus B. Leblanc, Llewellyn Green, Nandita R. Garud, Joshua M. Schmidt, Alexandre Fournier-Level, and Charles Robin. Structural variants and selective sweep foci contribute to insecticide resistance in the Drosophila genetic reference panel. G3: Genes, Genomes, Genetics, 8(11):3489–3497, November 2018. doi: 10.1534/g3.118.200619.

29. Belinda van Heerwaarden, Vanessa Kellermann, and Carla M. Sgrò. Limited scope for plasticity to increase upper thermal limits. Functional Ecology, 30(12):1947–1956, 2016. doi: 10.1111/1365-2435.12687.

30. Ben L. Phillips, Martha M. Muñoz, Amberlee Hatcher, Stewart L. Macdonald, John Llewelyn, Vanessa Lucy, and Craig Moritz. Heat hardening in a tropical lizard: Geographic variation explained by the predictability and variance in environmental temperatures. Functional Ecology, 30(7):1161–1168, 2016. doi: 10.1111/1365-2435.12609.

31. Suegene Noh, Elizabeth R. Everman, Christopher M. Berger, and Theodore J. Morgan. Seasonal variation in basal and plastic cold tolerance: Adaptation is influenced by both long-and short-term phenotypic plasticity. Ecology and Evolution, 7(14):5248–5257, 2017. doi: 10.1002/ece3.3112.

32. Rachael Morgan, Mette H. Finnøen, and Fredrik Jutfelt. CT max is repeatable and doesn’t reduce growth in zebrafish. Scientific Reports, 8(1):7099, 2018. doi: 10.1038/s41598-018-25593-4.

33. Anthony L. Gilbert and Donald B. Miles. Antagonistic responses of exposure to sublethal temperatures: Adaptive phenotypic plasticity coincides with a reduction in organismal performance. The American Naturalist, 194(3):344–355, 2019. doi: 10.1086/704208.

34. Andrew D. Nguyen, Megan Brown, Jordan Zitnay, Sara Helms Cahan, Nicholas J. Gotelli, Amy Arnett, and Aaron M. Ellison. Trade-offs in cold resistance at the northern range edge of the common woodland ant aphaenogaster picea (formicidae). The American Naturalist, 194(6):E151–E163, 2019. doi: 10.1086/705939.

35. Matthew C. Sasaki and Hans G. Dam. Integrating patterns of thermal tolerance and phenotypic plasticity with population genetics to improve understanding of vulnerability to warming in a widespread copepod. Global Change Biology, 25(12):4147–4164, 2019. doi: 10.1111/gcb.14811.

36. Matthew C. Sasaki and Hans G. Dam. Genetic differentiation underlies seasonal variation in thermal tolerance, body size, and plasticity in a short-lived copepod. Ecology and Evolution, 10(21):12200–12210, 2020. doi: 10.1002/ece3.6851.

37. Sean W. Deery, Julie E. Rej, Daniel Haro, and Alex. R. Gunderson. Heat hardening in a pair of anolis lizards: constraints, dynamics and ecological consequences. Journal of Experimental Biology, 224(7):jeb240994, 04 2021. ISSN 0022-0949. doi: 10.1242/jeb.240994.

38. Matthew C. Sasaki and Hans G. Dam. Negative relationship between thermal tolerance and plasticity in tolerance emerges during experimental evolution in a widespread marine invertebrate. Evolutionary Applications, 14(8):2114–2123, 2021. doi: 10.1111/eva.13270.

39. Erica O’Neill, Hannah E. Davis, and Heath A. MacMillan. A lack of repeatability creates the illusion of a trade-off between basal and plastic cold tolerance. Proceedings of the Royal Society B: Biological Sciences, 288(1964):20212121, 12 2021. ISSN 0962-8452. doi: 10.1098/rspb.2021.2121.

40. Giovanna Mottola, María E. López, Anti Vasemägi, Mikko Nikinmaa, and Katja Anttila. Are you ready for the heat? phenotypic plasticity versus adaptation of heat tolerance in three-spined stickleback. Ecosphere, 13(4):e4015, 2022. doi: 10.1002/ecs2.4015.

41. Yimen G. Araya-Ajoy and Niels J. Dingemanse. Repeatability, heritability, and age-dependence of seasonal plasticity in aggressiveness in a wild passerine bird. Journal of Animal Ecology, 2017. doi: 10.1111/1365-2656.12621.

42. Jolle W. Jolles, Helen D. Briggs, Yimen G. Araya-Ajoy, and Neeltje J. Boogert. Personality, plasticity and predictability in sticklebacks: Bold fish are less plastic and more predictable than shy fish. Animal Behaviour, 154:193–202, 2019. doi: 10.1016/j.anbehav.2019.06.022.

43. Marc T. J. Johnson and Zain Nassrullah. The improbability of detecting trade-offs and some practical solutions. Journal of Evolutionary Biology, 37(10):1205–1214, 2024. doi: 10.1093/jeb/voae096.

44. Juhan Javoiš. Ghost trade-offs: negative correlations between beneficial traits may be caused by other mechanisms than trade-offs. Theory in Biosciences, 145(3):34, 2026. doi: 10.1007/s12064-026-00471-0.

45. Douglas Bates, Reinhold Kliegl, Shravan Vasishth, and Harald Baayen. Parsimonious mixed models. arXiv preprint 1506.04967, 2015.

46. Wei Wang. Identifiability of covariance parameters in linear mixed effects models. Linear Algebra and its Applications, 506:603–613, 2016. doi: 10.1016/j.laa.2016.06.022.

47. R Core Team. R: A Language and Environment for Statistical Computing. R Foundation for Statistical Computing, Vienna, Austria, 2026.

48. Julien G. A. Martin, Daniel H. Nussey, Alastair J. Wilson, and Denis Réale. Measuring individual differences in reaction norms in field and experimental studies: A power analysis of random regression models. Methods in Ecology and Evolution, 2(4):362–374, 2011. doi: 10.1111/j.2041-210X.2010.00084.x.

49. Martijn van de Pol. Quantifying individual variation in reaction norms: How study design affects the accuracy, precision and power of random regression models. Methods in Ecology and Evolution, 3(2):268–280, 2012. doi: 10.1111/j.2041-210X.2011.00160.x.

50. Giovanni Bellu, Maria Pia Saccomani, Sandrine Audoly, and Luca D’Angiò. Daisy: Differential algebra for identifiability of systems, 2026. Software for structural identifiability analysis of ordinary differential equation models. Accessed 2026-09-28.

51. Thomas M. Houslay and Alastair J. Wilson. Avoiding the misuse of blup in behavioural ecology. Behavioral Ecology, 28(4):948–952, 2017. doi: 10.1093/beheco/arx023.

52. Niels J. Dingemanse and Ned A. Dochtermann. Quantifying individual variation in behaviour: mixed-effect modelling approaches. Journal of Animal Ecology, 82:39–54, 2013. doi: 10.1111/1365-2656.12013.

53. Richard Gomulkiewicz and Mark Kirkpatrick. Quantitative genetics and the evolution of reaction norms. Evolution, 46(2):390–411, 1992. doi: 10.1111/j.1558-5646.1992.tb02047.x.

54. Joel G. Kingsolver, Richard Gomulkiewicz, and Patrick A. Carter. Variation, selection and evolution of function-valued traits. Genetica, 112–113(1):87–104, 2001. doi: 10.1023/A:1013323318612.

55. Michael L. Logan and Christian L. Cox. Genetic constraints, transcriptome plasticity, and the evolutionary response to climate change. Frontiers in Genetics, 11:538226, 2020. doi: 10.3389/fgene.2020.538226.

56. Marta A. Santos, Marta A. Antunes, Afonso Grandela, Ana S. Quina, Mauro Santos, Margarida Matos, and Pedro Simões. Slow and population specific evolutionary response to a warming environment. Scientific Reports, 13:9700, 2023. doi: 10.1038/s41598-023-36273-3.

57. Lisa Bjerregaard Jørgensen, Hans Malte, Michael Ørsted, Nikolaj Andreasen Klahn, and Johannes Overgaard. A unifying model to estimate thermal tolerance limits in ectotherms across static, dynamic and fluctuating exposures to thermal stress. Scientific Reports, 11 (1):12840, 2021. doi: 10.1038/s41598-021-92004-6.

58. Félix Leiva, Mauro Santos, Edwin Niklitschek, Enrico Rezende, and Wilco Verberk. Genetic variation of heat tolerance in a model ectotherm: an approach using thermal death time curves. ecoevorxiv, 06 2024. doi: 10.32942/X20C9T.

59. Daniel H. Nussey, Alastair J. Wilson, and Jon E. Brommer. The evolutionary ecology of individual phenotypic plasticity in wild populations. Journal of Evolutionary Biology, 20:831–844, 2007.

60. Yaamini R. Venkataraman, Julia C. Kelso, Catlin Payne, Heidi L. Freitas, Jasmine Kohler, and Carolyn K. Tepolt. Plasticity, not genetics, shapes individual responses to thermal stress in non-native populations of the european green crab (Carcinus maenas). Integrative and Comparative Biology, 65(4):1148–1165, 2025. doi: 10.1093/icb/icaf131.

61. F. E. J. Fry, J. S. Hart, and K. F. Walker. Lethal temperature relations for a sample of young speckled trout, salvelinus fontinalis. Technical Report 66, Publications of the Ontario Fisheries Research Laboratory, 1946.

62. G. W. Bullard, L. B. Buckley, and J. G. Kingsolver. Incorporating variation in death times improves predictions of ectotherm responses to stressful temperatures. PLOS Biology, 24 (5):e3003623, 2026. doi: 10.1371/journal.pbio.3003623.

63. Daniel W. A. Noble, Pieter A. Arnold, Shinichi Nakagawa, and Patrice Pottier. A flexible modelling framework for estimating thermal tolerance and sensitivity. bioRxiv, 2026. doi: 10.64898/2026.07.16.738378.

64. Michael Ørsted, Lisa Bjerregaard Jørgensen, Petr Håla, and Johannes Overgaard. Integrating physiological rates of thermal stress and repair predicts heat failure during temperature fluctuations. Ecology Letters, 29(5):e70398, 2026. doi: 10.1111/ele.70398.

65. Douglas Bates, Martin Mächler, Ben Bolker, and Steve Walker. Fitting linear mixed-effects models using lme4. Journal of Statistical Software, 67(1):1–48, 2015. doi: 10.18637/jss.v067.i01.

66. Vincent Gardeux, Roel P. J. Bevers, Fabrice P. A. David, Emily Rosschaert, Romain Rochepeau, and Bart Deplancke. Dgrpool, a web tool leveraging harmonized drosophila genetic reference panel phenotyping data for the study of complex traits. eLife, 12:RP88981, 2024. doi: 10.7554/eLife.88981.

