## Supporting Information for "Separating biological signal from statistical artifact in trait trade-off inference"

Associated with the text *Separating biological signal from statistical artifact in trait trade-off inference*

October 2, 2026

#### Contents

**1 Supporting Figures and Tables** **2**

**2 Supporting Text** **13**

2.1 Analytical derivations of Table S3 formulas . . . . . 13

2.2 Estimation of the independent-parameter null model . . . . . 16

### 1 Supporting Figures and Tables

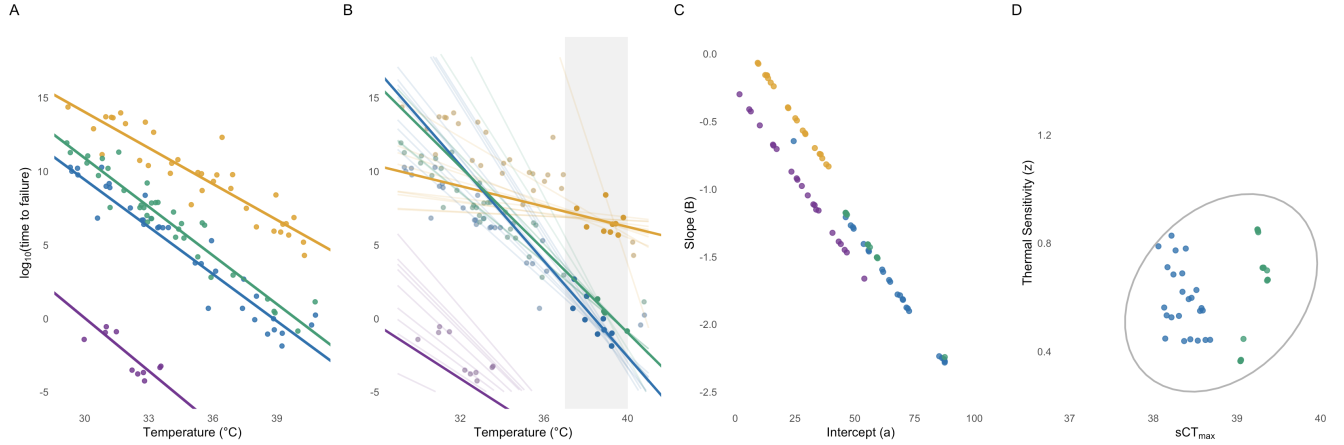


Figure S1: **Panel A:** Many sampled data points from 4 linear generating distributions with intercepts equal to those from Figure 2A, but slopes varying randomly between -0.7 and 1.3. **Panel B:** Estimated least squares fits (bold pink lines) based data within a restricted predictor range (gray shaded region). Light lines denote similarly likely fits obtained by bootstrap sampling the observed data of the same color(with replacement) and refitting the line. **Panel C:** Slope/intercept pairs for lines from panel B form a spurious correlation. **Panel D:** Reparameterization to z/sCTmax results in a muddy correlation, as slope variation was outside the normal range of single population TDT data.

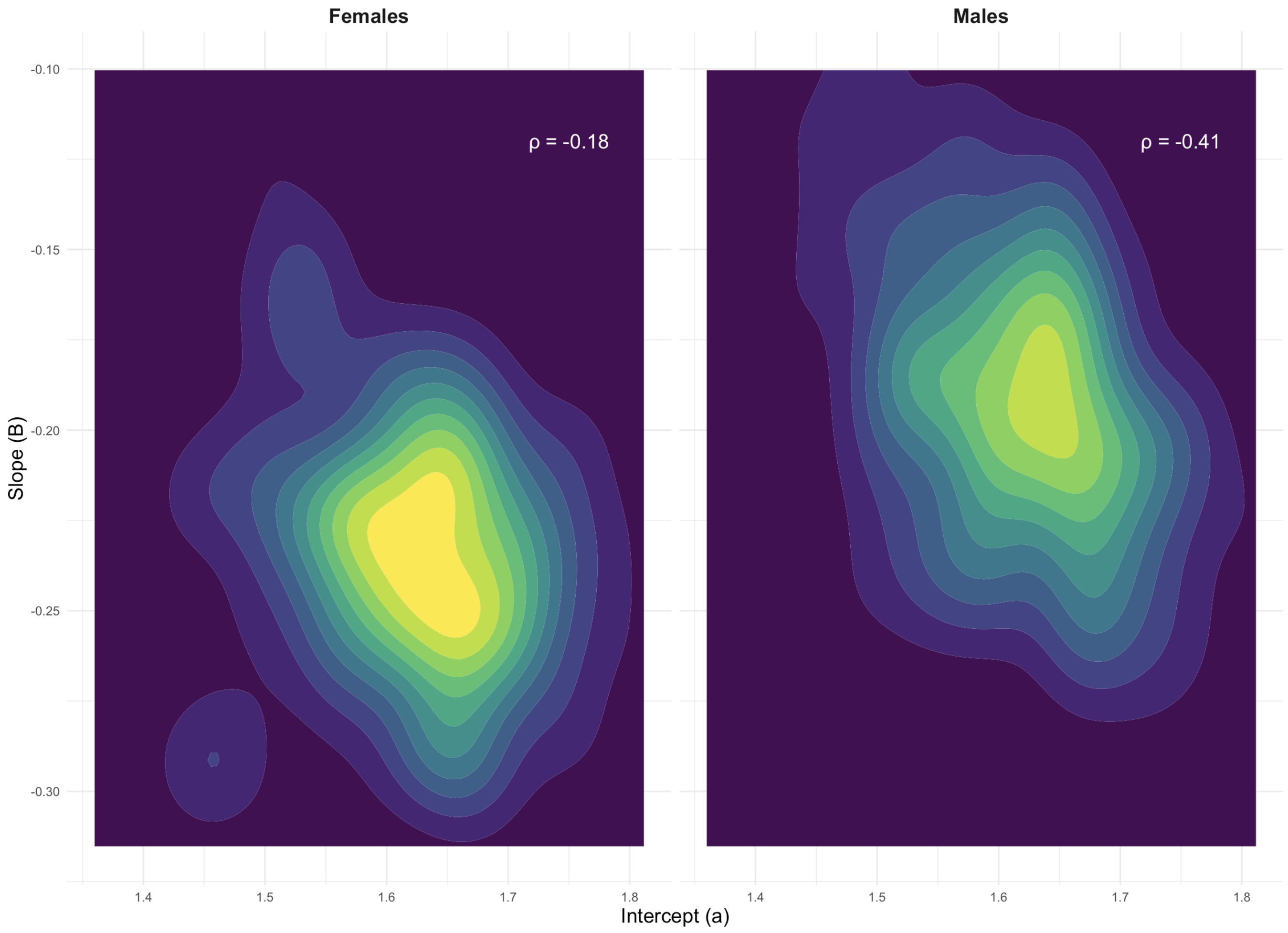


Figure S2: Joint density plot of Least-Squares estimated slope-intercept pairs for 100 isogenic lines of *D. melanogaster* after predictor centering. **Panel A:** Female data. **Panel B:** Male data.

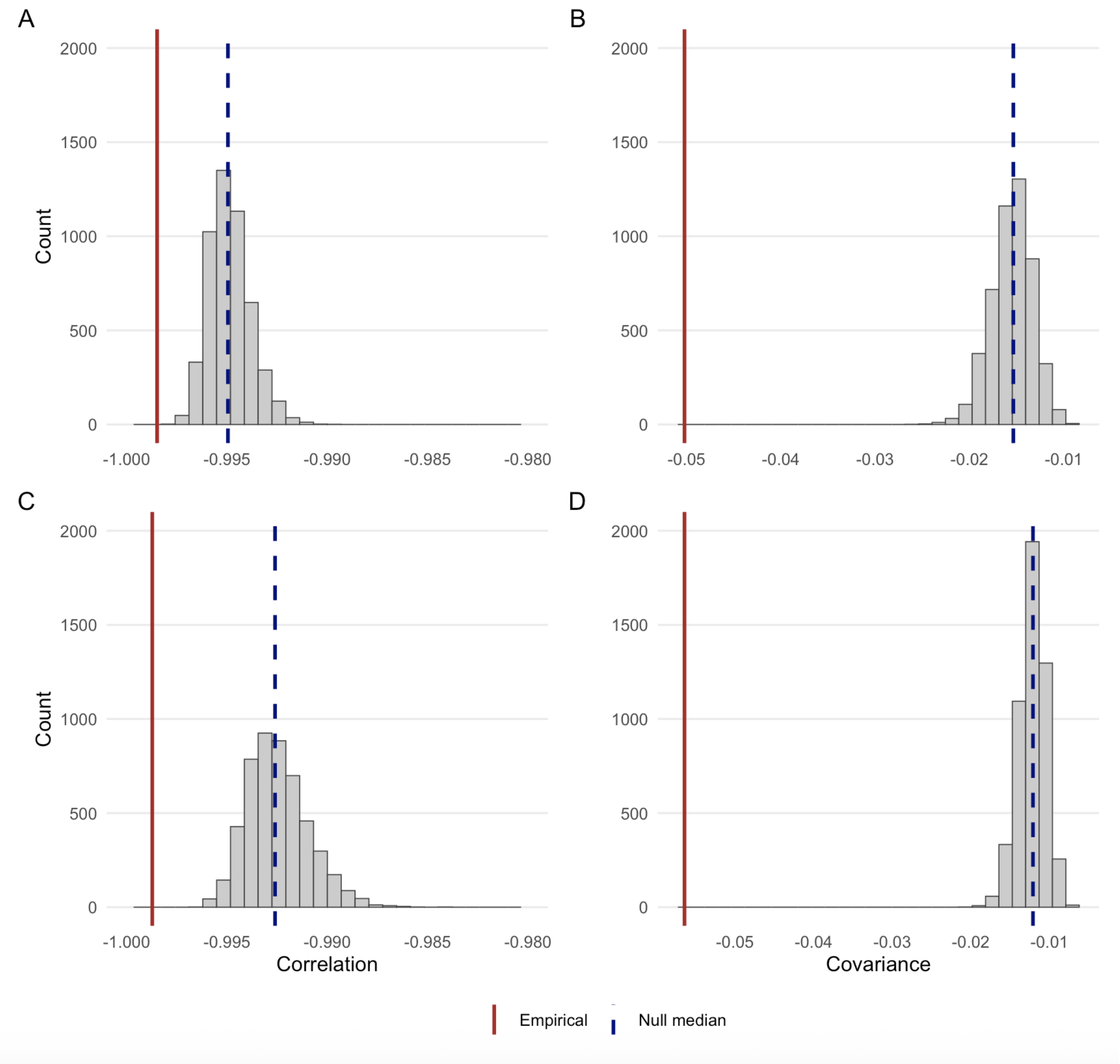


Figure S3: Results of our non-identifiability framework for the correlation (first column) and covariance (second column) between TDT slope and intercept for 100 female (first row) and male (second row) isogenic lines of *D. melanogaster*. The gray histogram displays the distribution of the statistic of interest expected under the assumption of no true covariation between these parameters, given the observed sampling procedure. The dashed blue line is the median of this null distribution, while the solid red line is the empirically observed value of the statistic of interest.

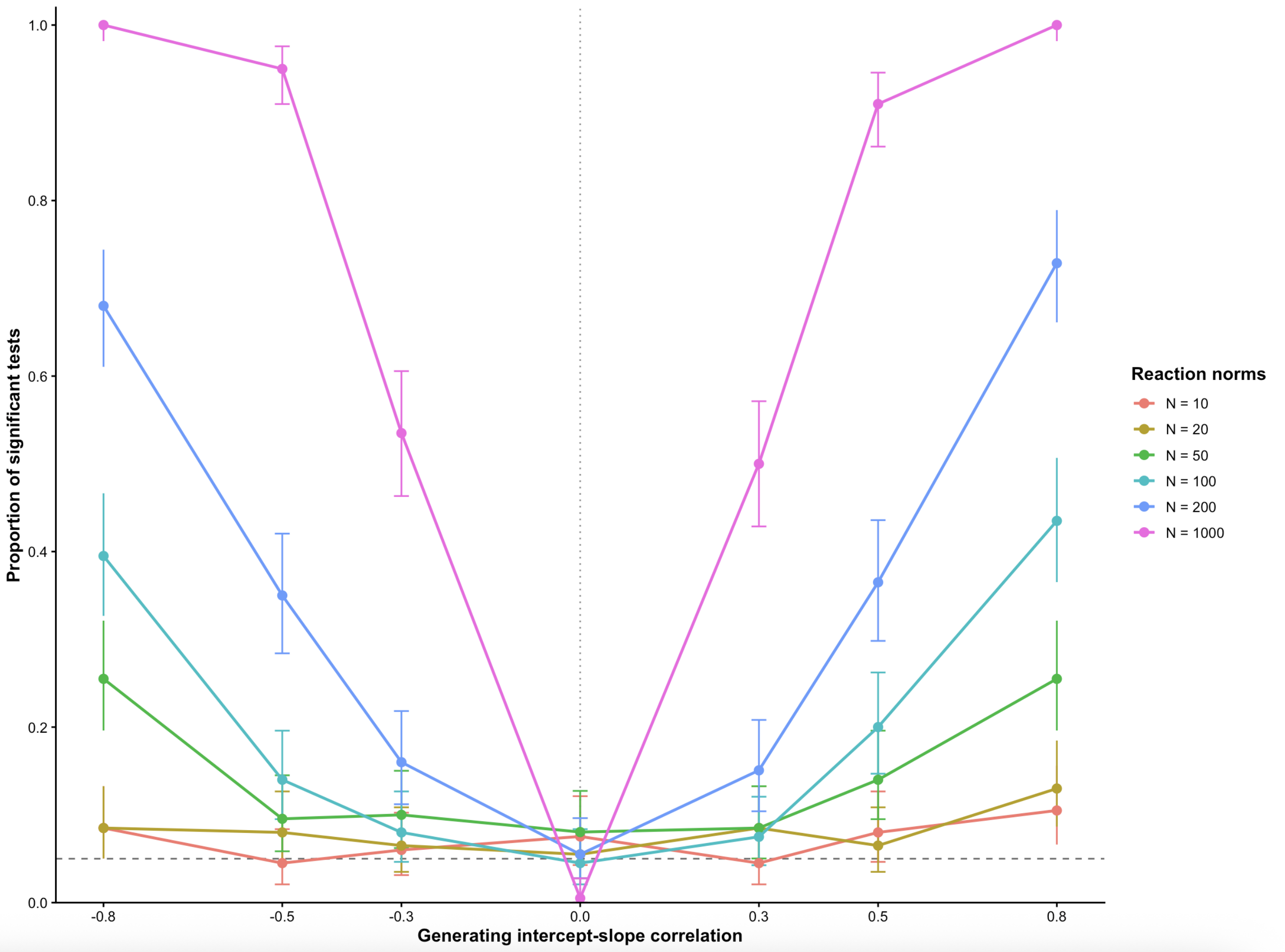


Figure S4: Power analysis results based on the isogenic *Drosophila* data. We assumed a generating slope mean and variance of -0.21 and 0.0018, and an intercept mean and variance of 1.6 and 0.006, respectively, according to the least squares estimates of these parameters. These are considered conservative upper bounds of possible variance estimates (due to non-identifiability-induced variance), and unbiased mean estimates. We drew 10 samples from each predictor (-1.5, -0.5, 0.5, 1.5), which is a centered version of the empirical protocol. Line colors denote the number of TDT curves used in the test.

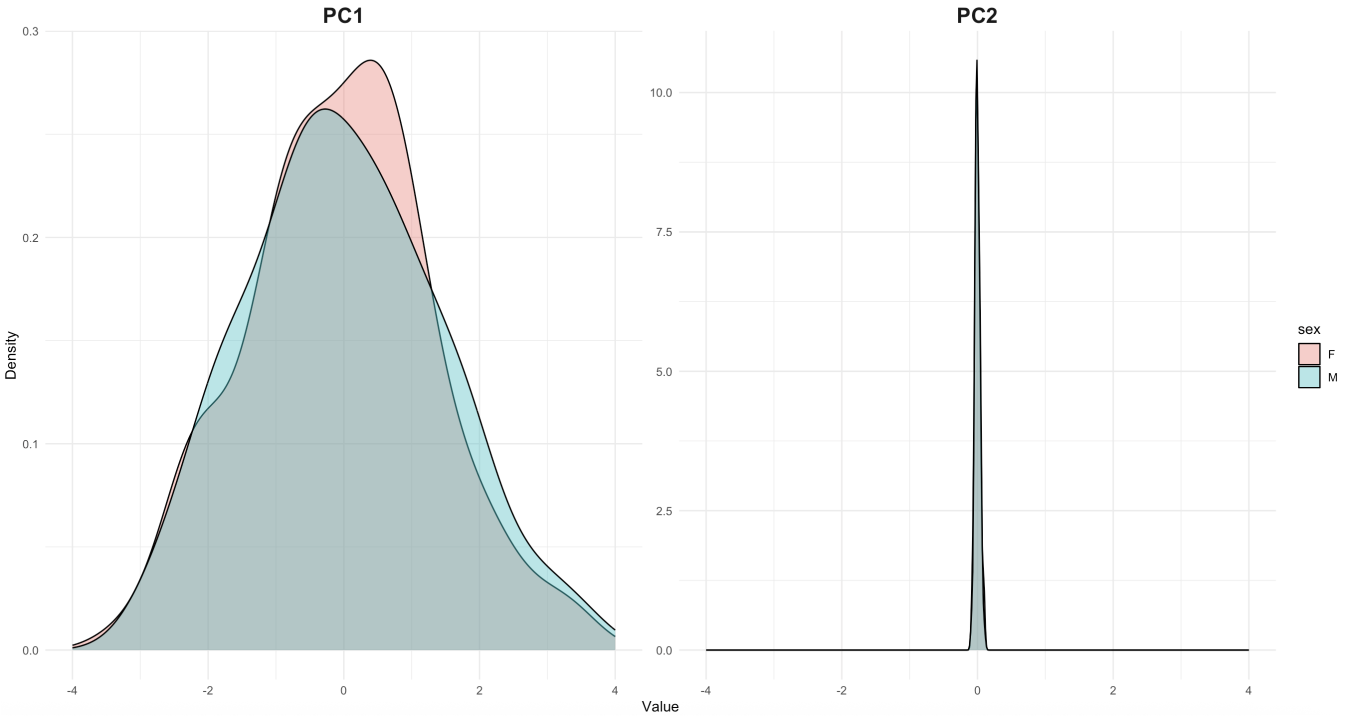


Figure S5: Density plots for PC1 and PC2 obtained from the joint distribution of estimated TDT slope and intercept for 100 isogenic populations of *D. melanogaster*.

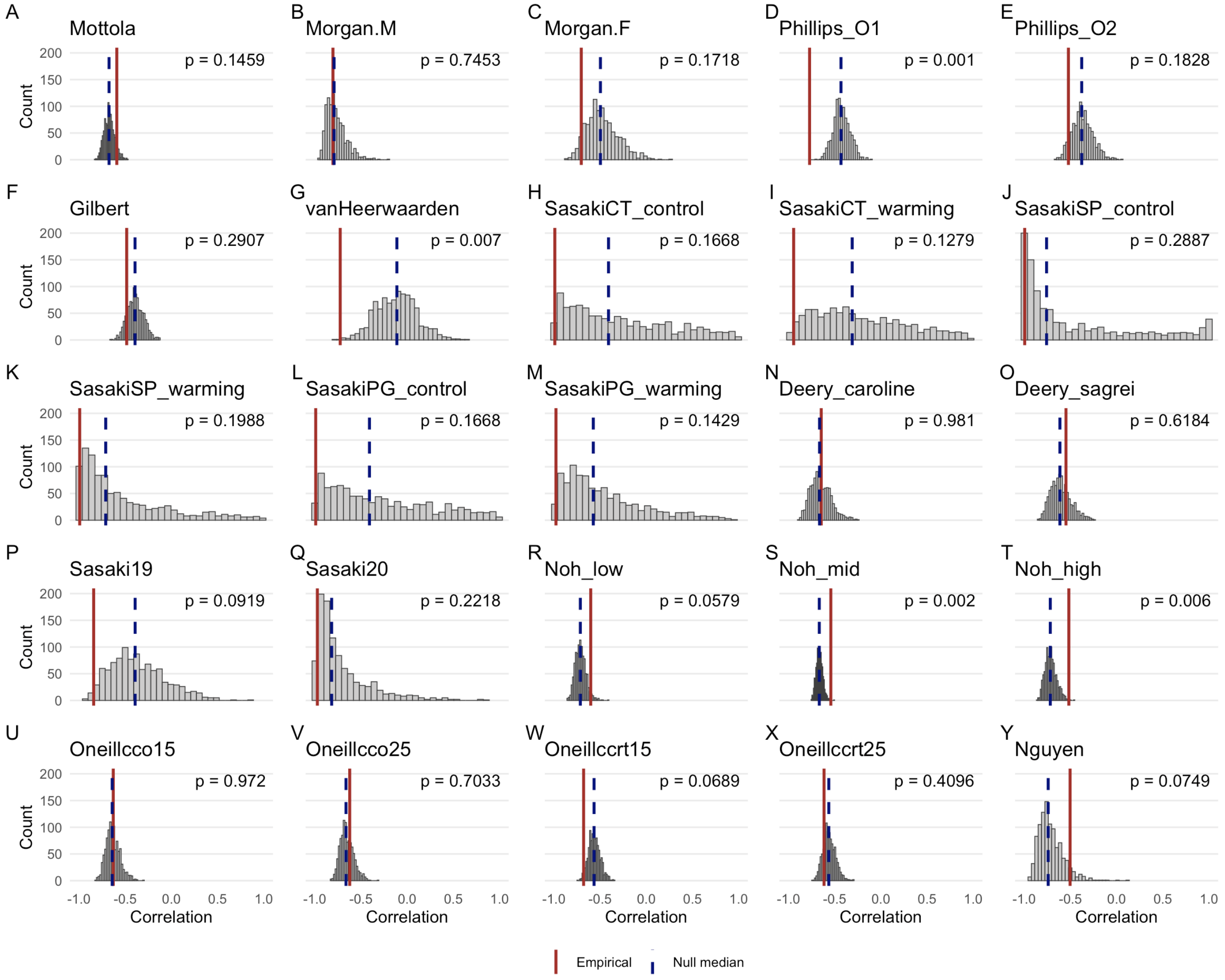


Figure S6: Bootstrapped distributions for the correlation between an initial thermal tolerance measurement (intercept) and an associated plasticity measurement (slope) expected if statistical artifact and sampling error were the sole cause of covariation between these parameters for 25 thermal tolerance reaction norm datasets. The gray histogram displays the distribution of the statistic of interest expected under the assumption of no true covariation between these parameters, given the observed sampling procedure. The dashed blue line is the median of this null distribution, while the solid red line is the empirically observed value of the statistic of interest. See [1] for more information on datasets and sources. Data sourced from [2–12].

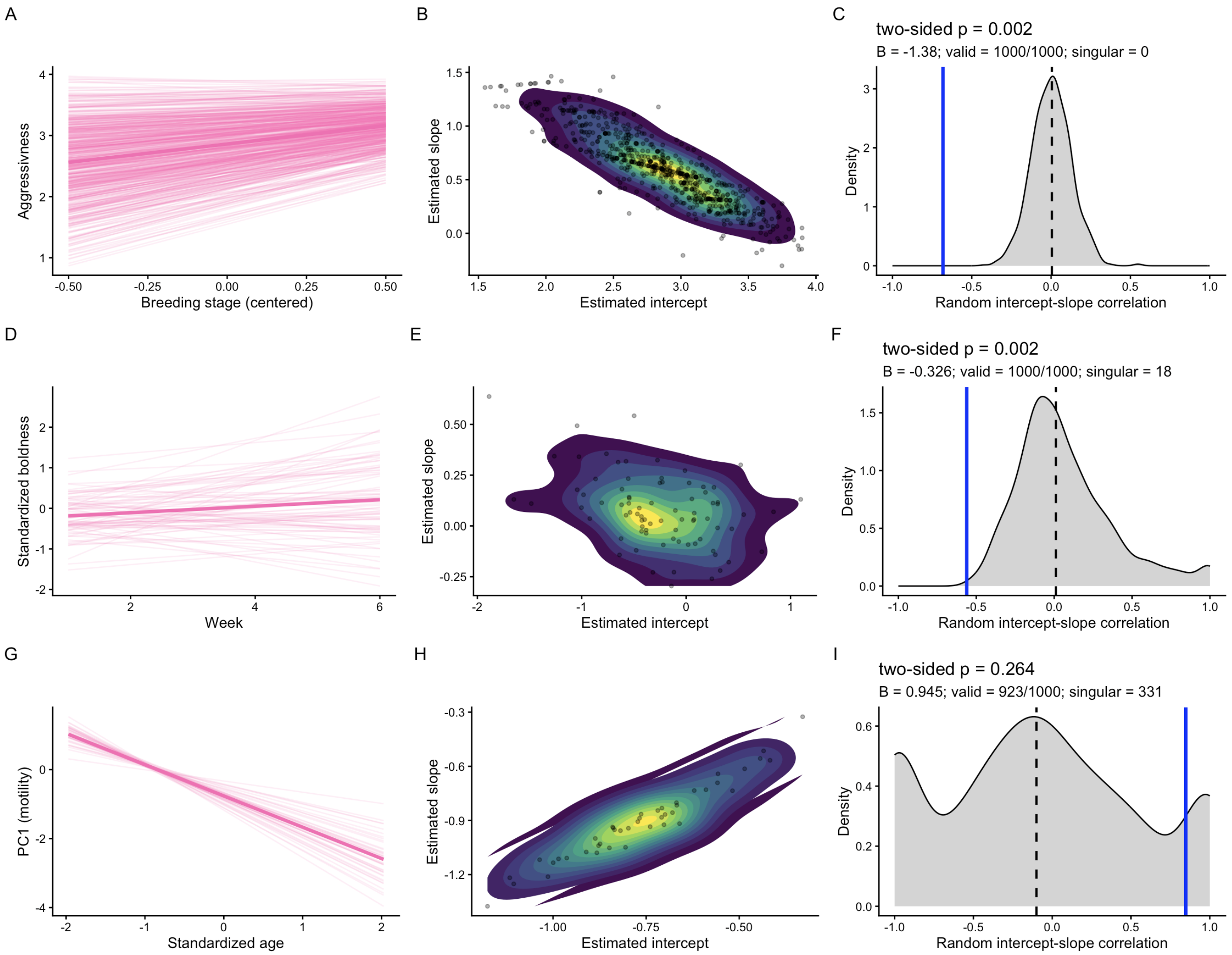


Figure S7: Reaction norms (column 1), joint distribution of slope and intercept estimates (column 2) and non-identifiability test results (column 3) for three literature reaction norm datasets investigating the trade-offs between male Great Tit (*Parus major*, aggressiveness and plasticity across breeding stages (row 1), Three-Spined Stickleback *Gasterosteus aculeatus* boldness and plasticity across time in a new environment (row 2), and between sperm motility and longevity in the marine mussel *Mytilus galloprovincialis* (row 3). All three datasets were modeled using linear mixed models, where displayed response variables were fixed effects. In the first column, thin lines represent reaction norm estimates for each given group, while the thick line represents the best-fit line for the population as a whole. In the second row, yellow tones represent regions of high density, blue tones represent regions of lower density. In the third row, the gray distribution displays the distribution of the statistic of interest expected under the assumption of no true covariation between these parameters, given the observed sampling procedure, while the blue line displays the emperically observed correlation. Data sourced from [**https://doi.org/10.1002/ece3.8514**, 13, 14].

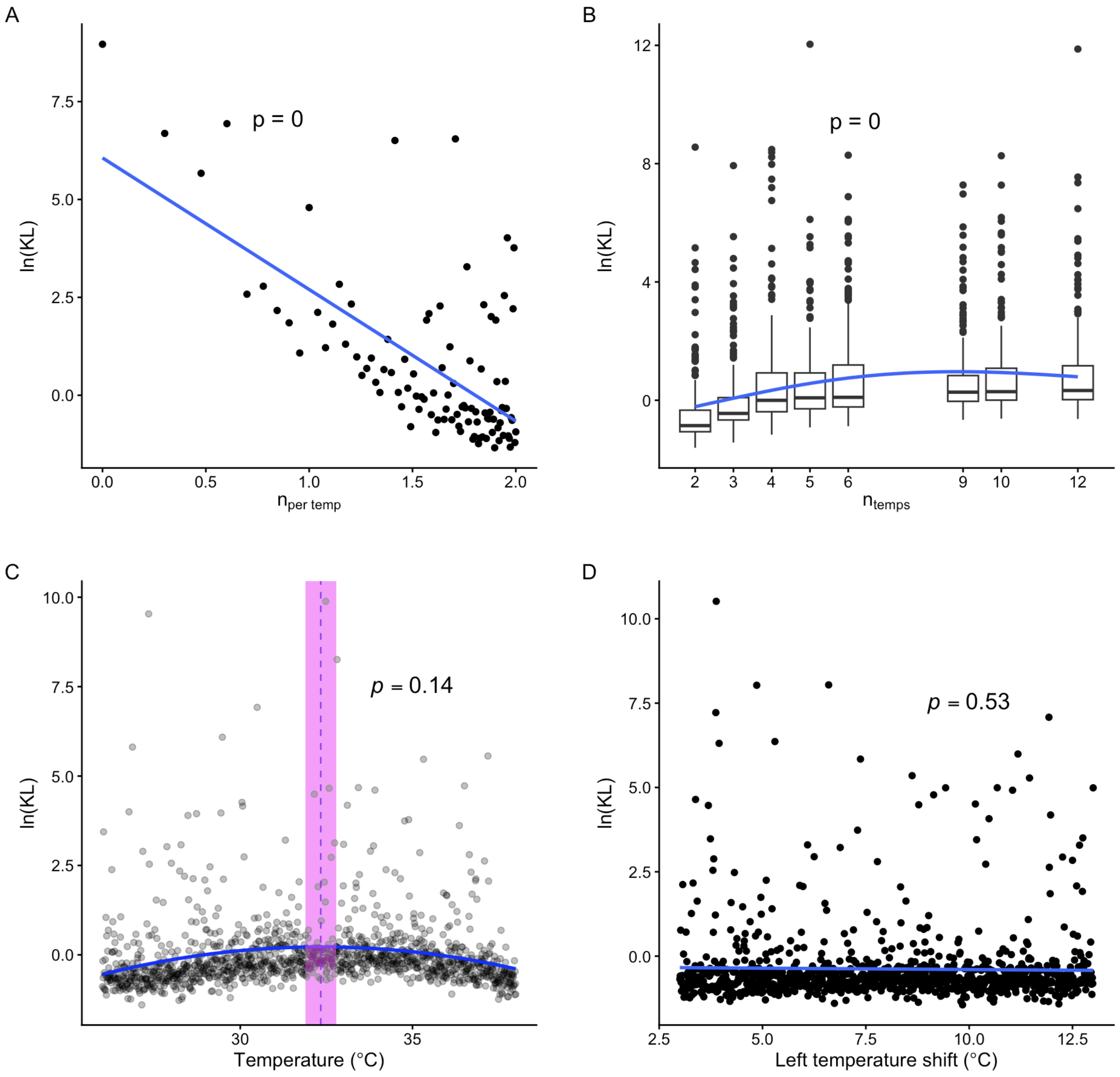


Figure S8: log KL divergence for known and estimated joint distributions of $z/sCT_{max}$ from TDT data simulated from a known generating distribution under various sampling conditions. For each data point, 100 TDT curves were drawn from a known distribution of $z$ and $sCT_{max}$. For each of these curves, $n_{per\ temp}$ failure times were simulated at $n_{temps}$ different temperatures. The generating TDT distribution is as follows: $(z, sCT_{\text{max}}) \sim MVN(\mu, \Sigma)$, $Cov(z, sCT_{\text{max}}) = 0$, $\sigma_z = 1\ min^{-1}$, $\sigma_{sCT_{\text{max}}} = 1\ °\text{C} \cdot min^{-1}, \mu_z = 4.5\ min^{-1}, \mu_{\text{sCTmax}} = 46\ °\text{C} \cdot min^{-1}, \sigma^2_{\varepsilon} = 0.01.$, where means were based on empirical data for *D. melanogaster*, while variance parameters were arbitrarily set to 1, as realistic ranges of these parameters are unknown due to sampling error. However, adjusting these parameters yields similar results (see supplemental code). $n_{per\ temp} = 15$ unless otherwise noted. **Panel A**: the effect of replication per temperature per line ($n_{per\ temp}$) on reestimation quality. Here, $n_{temps} = 5$, and spanned from 28-36C. The blue line is ordinary least squares regression, and the p-value is associated with the slope of this line. **Panel B**: The effect of $n_{temps}$ on reestimation quality. $n_{temps}$ equally spaced temperatures were used within the range 28-38C. **Panel C**: Effect of intermediate temperature placement on reestimation quality. Three temperatures were used: two fixed temperatures at 26C and 38C, and an intermediate temperature that ranged incrementally between those values. The quadratic fit indicates how model fit changed with the choice of this intermediate temperature, and the vertex of this parabola (pink vertical line) is displayed along with a 95% confidence interval (shaded pink region). The p-value is associated with the null hypothesis that the midpoint of the temperature range is equal to the vertex. **Panel D**: Effect of shifting the temperature window relative to $sCT_{max}$: three equally spaced temperatures were placed at many different locations, and ln(KL) divergence is shown relative to the difference between $sCT_{max}$ and the maximum temperature sampled. The p-value tests whether the slope of the least-squares fit (blue line) differs from 0.

Table S1: GWAS results for 4 parameter-phenotypes. PC1 and PC2 are in $z/sCT_{max}$ parameter space. Each cell contains the number of SNPs significant to the $10^{-5}$ level (DGRP standard (Battlay et al 2018)) for both the column and row parameters, for females/males, respectively. The diagonal contains the number of significant SNPs for a given parameter. $\sim$20 significant SNPs are expected by chance ($\sim$2,000,000 SNPs $*10^{-5}$).

| | PC1 | PC2 | $z$ | $sCT_{max}$ |
|---|---|---|---|---|
| PC1 | 132/28 | 0/0 | 81/25 | 101/10 |
| PC2 | — | 15/28 | 0/0 | 0/0 |
| $\beta$ | — | — | 98/27 | 62/11 |
| $sCT_{max}$ | — | — | — | 122/14 |

Table S2: Correlation analysis results for raw empirical correlation (6th column), correction for regression to the mean (RTM) (7th column), and our identifiability test (8th column). The comparison column reports ”greater” if the empirical correlation is greater than the median null correlation expected in the absence of biological (in value, not magnitude), and ”less” if the reverse is true (see Figure S6 for full results). Other table contents and RTM results sourced from [1]. Bold values indicate significant differences from 0 to the $p < 0.05$ level (no adjustment for multiple tests was made for consistency with the analyzed literature). Entries above the horizontal line studied heat tolerance, studies below analyzed cold tolerance.

| Taxon | Species | Level | Trait | Treatment | Empirical $r$ | Null $r$ | Comparison |
|---|---|---|---|---|---|---|---|
| Fish | *Gasterosteus aculeatus* | Individual | CTmax | — | **-0.59** | 0.02 | Greater |
| Fish | *Danio rerio* | Individual | CTmax | Females | **-0.70** | -0.20 | Less |
| Fish | *D. rerio* | Individual | CTmax | Males | **-0.80** | 0.02 | Less |
| Reptile | *Anolis sagrei* | Individual | CTmax | — | **-0.55** | 0.04 | Greater |
| Reptile | *A. carolinensis* | Individual | CTmax | — | **-0.65** | 0.00 | Greater |
| Reptile | *Lampropholis coggeri* | Individual | CTmax | Observer 1 | **-0.77** | **-0.49** | **Less** |
| Reptile | *L. coggeri* | Individual | CTmax | Observer 2 | **-0.52** | -0.14 | Less |
| Reptile | *Urosaurus ornatus* | Individual | CTmax | — | **-0.48** | -0.09 | Less |
| Insect | *Drosophila melanogaster* | Population | CTmax | — | **-0.73** | **-0.65** | **Less** |
| Crustacean | *Acartia tonsa* | Population | LD50 | — | **-0.84** | -0.57 | Less |
| Crustacean | *A. tonsa* | Population | LD50 | — | **-0.97** | 0.66 | Less |
| Crustacean | *A. tonsa* | Selection line | LD50 | CT pop., control | **-0.99** | **-0.97** | **Less** |
| Crustacean | *A. tonsa* | Selection line | LD50 | CT pop., warming | **-0.94** | **-0.89** | Less |
| Crustacean | *A. tonsa* | Selection line | LD50 | SP pop., control | **-1.00** | -0.93 | Less |
| Crustacean | *A. tonsa* | Selection line | LD50 | SP pop., warming | **-0.99** | **-0.99** | Less |
| Crustacean | *A. tonsa* | Selection line | LD50 | PG pop., control | **-0.97** | -0.91 | **Less** |
| Crustacean | *A. tonsa* | Selection line | LD50 | PG pop., warming | **-0.98** | **-0.97** | Less |
| Insect | *D. melanogaster* | Female lines | CCRT | Low heat | **-0.81** | 0.01 | Greater |
| Insect | *D. melanogaster* | Female lines | CCRT | Mid heat | **-0.77** | -0.04 | **Greater** |
| Insect | *D. melanogaster* | Female lines | CCRT | High heat | **-0.87** | 0.02 | **Greater** |
| Insect | *D. melanogaster* | Individuals | CCRT | 15°C acclimated | **-0.68** | -0.09 | Less |
| Insect | *D. melanogaster* | Individuals | CCRT | 25°C acclimated | **-0.62** | -0.03 | Less |
| Insect | *D. melanogaster* | Individuals | CCO | 15°C acclimated | **-0.63** | 0.00 | Greater |
| Insect | *D. melanogaster* | Individuals | CCO | 25°C acclimated | **-0.62** | 0.01 | Greater |
| Insect | *Aphaenogaster picea* | Populations | CCRT | — | **-0.88** | 0.04 | Greater |

Table S3: A list of analytically derived effects of unidentifiability on TDT parameter estimates in terms of empirical quantities. We list whether the artifact is dictated by structural or practical non-identifiability.

| **Source of artifact** | **Relevant formula** | **Practical takeaway** |
|---|---|---|
| Structural | $\operatorname{cov}(\hat{\alpha}, \hat{\beta}) = \operatorname{cov}(\alpha, \beta) - \dfrac{\sigma_\varepsilon^2 \, \overline{T}}{n_{\text{temps}} \operatorname{var}(T)}$ | Centering data eliminates structural correlation/covariance bias. Note that this does not remove ridge structure. |
| Practical and structural | $\hat{\sigma}_\alpha^2 = \sigma_\alpha^2 + \dfrac{\sigma_\varepsilon^2}{n_{\text{total}}} \left(1 + \dfrac{T_{\text{bar}}^2}{\operatorname{var}(T)}\right)$ | Increasing overall sample size and limiting sampling error reduce intercept variance bias. Centering also reduces intercept bias. |
| Practical | $\hat{\sigma}_\beta^2 = \sigma_\beta^2 + \dfrac{\sigma_\varepsilon^2}{n_{\text{rep}} \operatorname{var}(T)}$ | Increasing per-temperature sample size, spreading sample temperatures across a wide range, and reducing sampling error reduce slope variance bias. |
| Practical | Ridge strength $\propto \dfrac{1}{\operatorname{var}(T)}$ | After centering, the magnitude of unidentifiability depends only on the spread of temperatures, not the number of samples. |
| Practical | $\operatorname{corr}(\hat{\alpha}, \hat{\beta})$ increases with $\dfrac{1}{\operatorname{var}(T)}$ | After centering, correlation bias depends only on the spread of temperatures, not the number of samples. |

### 2 Supporting Text

#### 2.1 Analytical derivations of Table S3 formulas

Below, we derive the relationships summarized in Table S3. We consider two distinct sources of artifact in linear trait estimation: geometric effects arising from uncentered predictors, and ridge geometry arising from correlated parameter values in similarly likely models.

These mechanisms affect parameter estimates differently and are governed by distinct properties of experimental design.

##### Model specification

$$\log(t_f) = \beta T + \alpha + \varepsilon, \quad \varepsilon \sim N(0, \sigma_\varepsilon^2), \quad (\alpha, \beta) \sim MVN(\mu, \Sigma)$$

##### Parameter definitions

- $T$: static exposure temperature
- $t_f$: time to failure under $T$
- $n_{\text{temps}}$: number of stressful temperatures sampled
- $n_{\text{per temp}}$: number of individuals sampled at each temperature
- $n_{\text{total}} = n_{\text{per temp}} \cdot n_{\text{temps}}$: total sample size
- $\bar{T}$: mean trial temperature
- $SS_T$: sum of squares of trial temperatures
- $\sigma_\alpha^2,\ \sigma_\beta^2,\ \sigma_\varepsilon^2$: variances of intercept, slope, and residual error
- $\hat{\alpha}, \hat{\beta}$: estimated intercept and slope for a given curve

##### Useful property

Under our assumed sampling conditions:

$$SS_T = \sum_{i=1}^{n_{\text{total}}} (T_i - \bar{T})^2 = \sum_{j=1}^{n_{\text{temps}}} n_{\text{per temp}} (T_j - \bar{T})^2 = n_{\text{per temp}} \sum_{j=1}^{n_{\text{temps}}} (T_j - \bar{T})^2$$

$$\text{Var}(T) = \frac{1}{n_{\text{temps}}} \sum_{j=1}^{n_{\text{temps}}} (T_j - \bar{T})^2$$

$$\Rightarrow \sum_{j=1}^{n_{\text{temps}}} (T_j - \bar{T})^2 = n_{\text{temps}} \cdot \text{Var}(T)$$

$$\Rightarrow SS_T = n_{\text{per temp}} \cdot n_{\text{temps}} \cdot \text{Var}(T) = n_{\text{total}} \cdot \text{Var}(T)$$

We use this equation throughout.

##### Intercept variance

From standard linear model theory ([15]):

$$\text{Var}(\hat{\alpha}) = \sigma_\alpha^2 + \sigma_\varepsilon^2 \left( \frac{1}{n_{\text{total}}} + \frac{\bar{T}^2}{SS_T} \right)$$

Substituting $SS_T$:

$$\text{Var}(\hat{\alpha}) = \sigma_\alpha^2 + \sigma_\varepsilon^2 \left( \frac{1}{n_{\text{total}}} + \frac{\bar{T}^2}{n_{\text{total}} \cdot \text{Var}(T)} \right)$$

The second term depends on $\bar{T}$ and can be eliminated by centering predictors. After centering:

$$\boxed{\mathrm{Var}(\hat{\alpha}) = \sigma_\alpha^2 + \frac{\sigma_\varepsilon^2}{n_{\text{total}}}}$$

Practical reduction of bias in intercept variance (unsurprisingly) relies on increasing total sample size or decreasing sampling error.

##### Slope variance

([15])

$$\mathrm{Var}(\hat{\beta}) = \sigma_\beta^2 + \frac{\sigma_\varepsilon^2}{SS_T} = \sigma_\beta^2 + \frac{\sigma_\varepsilon^2}{n_{\text{total}} \cdot \mathrm{Var}(T)}$$

Unlike the intercept, bias in slope variance does not depend on $\bar{T}$ and is therefore unaffected by centering. However, it decreases with both sample size and the spread of sampled temperatures.

—

##### Covariance derivation

For curve $j$:

$$\hat{\alpha}_j = \alpha_j + e_{\alpha j}, \quad \hat{\beta}_j = \beta_j + e_{\beta j}$$

Across curves:

$$\mathrm{Cov}(\hat{\alpha}, \hat{\beta}) = \mathrm{Cov}(\alpha_j + e_{\alpha j},\ \beta_j + e_{\beta j})$$

$$= \mathrm{Cov}(\alpha_j, \beta_j) + \mathrm{Cov}(\alpha_j, e_{\beta j}) + \mathrm{Cov}(e_{\alpha j}, \beta_j) + \mathrm{Cov}(e_{\alpha j}, e_{\beta j})$$

Assuming independence between parameters and estimation error ([15]):

$$\mathrm{Cov}(\hat{\alpha}, \hat{\beta}) = \mathrm{Cov}(\alpha, \beta) + \mathrm{Cov}(e_\alpha, e_\beta)$$

Using the standard least-squares result ([15]):

$$\mathrm{Cov}(e_\alpha, e_\beta) = -\frac{\sigma_\varepsilon^2 \bar{T}}{SS_T} = -\frac{\sigma_\varepsilon^2 \bar{T}}{n_{\text{total}} \cdot \mathrm{Var}(T)}$$

$$\boxed{\mathrm{Cov}(\hat{\alpha}, \hat{\beta}) = \mathrm{Cov}(\alpha, \beta) - \frac{\sigma_\varepsilon^2 \bar{T}}{n_{\text{total}} \cdot \mathrm{Var}(T)}}$$

Thus, centering ($\bar{T} = 0$) removes geometry-induced bias in covariance.

##### Ridge geometry and induced correlation

Even after centering, parameter estimates exhibit structured uncertainty due to the geometry of linear fitting.

Define the estimation covariance matrix:

$$\Sigma_e = \begin{pmatrix} \mathrm{Var}(e_\alpha) & \mathrm{Cov}(e_\alpha, e_\beta) \\ \mathrm{Cov}(e_\alpha, e_\beta) & \mathrm{Var}(e_\beta) \end{pmatrix}$$

Ridge geometry is determined by the eigenvalues $\lambda_1, \lambda_2$ of $\Sigma_e$, which define the principal axes of uncertainty (PC1 and PC2).

We define ridge strength as the ratio of variance along the principal directions:

$$\text{Ridge strength} = \frac{\lambda_1}{\lambda_2}, \quad \lambda_1 \geq \lambda_2$$

After centering, covariance vanishes, and:

$$\mathrm{Var}(e_\alpha) = \frac{\sigma_\varepsilon^2}{n_{\text{total}}}, \quad \mathrm{Var}(e_\beta) = \frac{\sigma_\varepsilon^2}{SS_T}$$

Thus:

$$\Sigma_e = \begin{pmatrix} \frac{\sigma_\varepsilon^2}{n_{\text{total}}} & 0 \\ 0 & \frac{\sigma_\varepsilon^2}{SS_T} \end{pmatrix}$$

Since $\Sigma_e$ is diagonal, its eigenvalues are:

$$\lambda_1 = \max\left(\frac{\sigma_\varepsilon^2}{n_{\text{total}}}, \frac{\sigma_\varepsilon^2}{SS_T}\right), \quad \lambda_2 = \min\left(\frac{\sigma_\varepsilon^2}{n_{\text{total}}}, \frac{\sigma_\varepsilon^2}{SS_T}\right)$$

Crucially, the ordering depends on $\text{Var}(T)$ through:

$$SS_T = n_{\text{total}} \cdot \text{Var}(T)$$

Substituting:

$$\text{Var}(e_\beta) = \frac{\sigma_\varepsilon^2}{n_{\text{total}} \cdot \text{Var}(T)}$$

For small $\text{Var}(T)$, we have:

$$\text{Var}(e_\beta) \gg \text{Var}(e_\alpha)$$

Thus:

$$\lambda_1 = \frac{\sigma_\varepsilon^2}{n_{\text{total}} \cdot \text{Var}(T)}, \quad \lambda_2 = \frac{\sigma_\varepsilon^2}{n_{\text{total}}}$$

And:

$$\text{Ridge strength} = \frac{\lambda_1}{\lambda_2} = \frac{\sigma_\varepsilon^2/(n_{\text{total}} \cdot \text{Var}(T))}{\sigma_\varepsilon^2/n_{\text{total}}}$$

Cancelling $\sigma_\varepsilon^2$:

$$= \frac{1}{\text{Var}(T)}$$

$$\boxed{\text{Ridge strength} \propto \frac{1}{\text{Var}(T)}}$$

Decreasing temperature variance elongates the estimation uncertainty along a single direction, producing a ridge of similarly likely parameter values. We assert proportionality rather than equality, because ”ridge strength” is a unitless quantity of arbitrary scaling.

**Induced correlation**

The observed correlation across curves is:

$$\text{Corr}(\hat{\alpha}, \hat{\beta}) = \frac{\text{Cov}(\hat{\alpha}, \hat{\beta})}{\sqrt{\text{Var}(\hat{\alpha})\text{Var}(\hat{\beta})}}$$

After centering, $\text{Cov}(e_\alpha, e_\beta) = 0$, so covariance is not directly induced within a single fit. However, across multiple curves, the distribution of estimates is governed by $\Sigma_e$.

When ridge strength is large ($\lambda_1 \gg \lambda_2$), variation across curves is dominated by a single principal direction:

$$(\hat{\alpha}, \hat{\beta}) \approx z\mathbf{v}_1$$

What this means is that when minimal variation in linear regression parameters is present, estimated parameters will lie approximately on this line, or ”ridge”, spaced by their respective values of $z$, which is determined by sampling and estimation error. So, in the absence of trait variation, and in the presence of low temperature variation, when ridge strength is high:

$$|\text{Corr}(\hat{\alpha}, \hat{\beta})| \to 1$$

Since ridge strength scales as $1/\text{Var}(T)$, we obtain:

$$\boxed{|\text{Corr}(\hat{\alpha}, \hat{\beta})| \text{ increases as } \frac{1}{\text{Var}(T)} \text{ increases}}$$

Thus, even in the absence of direct covariance, estimation uncertainty induces apparent correlation across curves.

#### 2.2 Estimation of the independent-parameter null model

We estimated a generative null model in which reaction-norm intercepts and slopes varied among groups but were independent at the latent level. For observation $i$ in group $g$, the model was

$$y_{ig} = (\beta_0 + a_g) + (\beta_1 + b_g)x_{ig} + \varepsilon_{ig}, \tag{1}$$

where $\beta_0$ and $\beta_1$ denote the population mean intercept and slope, respectively, and $a_g$ and $b_g$ denote group-specific deviations from these means. The group-level effects were assumed to follow

$$\begin{pmatrix} a_g \\ b_g \end{pmatrix} \sim \mathcal{N}\left[\begin{pmatrix} 0 \\ 0 \end{pmatrix}, \begin{pmatrix} \tau_a^2 & 0 \\ 0 & \tau_b^2 \end{pmatrix}\right]. \tag{2}$$

Thus, the latent intercept–slope covariance was fixed at zero, whereas the intercept variance $\tau_a^2$ and slope variance $\tau_b^2$ were estimated from the data. Observation-level errors were assumed to be independent and normally distributed,

$$\varepsilon_{ig} \sim \mathcal{N}(0, \sigma_e^2). \tag{3}$$

The predictor values were treated as fixed and were retained exactly as observed within each group. When predictor centering was used, $x_{ig}$ represented the observed predictor minus the specified centering value. Consequently, the intercept represented the expected response at that predictor value. For group $g$, let $\mathbf{y}_g$ be the vector of responses and let

$$\mathbf{X}_g = \begin{pmatrix} 1 & x_{1g} \\ \vdots & \vdots \\ 1 & x_{n_g g} \end{pmatrix} \tag{4}$$

be the corresponding design matrix. After integrating over the group-level random effects, the marginal distribution of the observations was

$$\mathbf{y}_g \sim \mathcal{N}\left(\mathbf{X}_g\boldsymbol{\beta}, \mathbf{V}_g\right), \tag{5}$$

where

$$\boldsymbol{\beta} = \begin{pmatrix} \beta_0 \\ \beta_1 \end{pmatrix}, \qquad \mathbf{V}_g = \mathbf{X}_g\mathbf{D}\mathbf{X}_g^{\mathsf{T}} + \sigma_e^2\mathbf{I}_{n_g}, \tag{6}$$

and

$$\mathbf{D} = \begin{pmatrix} \tau_a^2 & 0 \\ 0 & \tau_b^2 \end{pmatrix}. \tag{7}$$

The diagonal structure of $\mathbf{D}$ defines the independent-parameter null hypothesis,

$$H_0 : \mathrm{Cov}(a_g, b_g) = 0. \tag{8}$$

The model parameters were estimated by maximizing the full marginal likelihood, rather than a restricted likelihood. Assuming independence among groups, the negative log-likelihood minimized during optimization was

$$-\ell(\boldsymbol{\theta}) = \frac{1}{2}\sum_{g=1}^{G}\left[n_g \log(2\pi) + \log|\mathbf{V}_g| + (\mathbf{y}_g - \mathbf{X}_g\boldsymbol{\beta})^{\mathsf{T}}\,\mathbf{V}_g^{-1}\,(\mathbf{y}_g - \mathbf{X}_g\boldsymbol{\beta})\right], \tag{9}$$

where the free parameters were

$$\boldsymbol{\theta} = \left(\beta_0,\, \beta_1,\, \log\tau_a^2,\, \log\tau_b^2,\, \log\sigma_e^2\right). \tag{10}$$

Variance components were optimized on the logarithmic scale and exponentiated when evaluating the likelihood. This parameterization ensured that all variance estimates remained strictly positive during optimization. The likelihood was evaluated separately for each group using a Cholesky factorization of $\mathbf{V}_g$. If $\mathbf{R}_g$ denotes the upper-triangular Cholesky factor satisfying

$$\mathbf{R}_g^{\mathsf{T}}\mathbf{R}_g = \mathbf{V}_g, \tag{11}$$

then the log determinant was calculated as

$$\log|\mathbf{V}_g| = 2\sum_j \log R_{g,jj}. \tag{12}$$

The quadratic term was calculated by solving

$$\mathbf{R}_g^{\mathsf{T}}\mathbf{z}_g = \mathbf{y}_g - \mathbf{X}_g\boldsymbol{\beta} \tag{13}$$

and evaluating $\mathbf{z}_g^{\mathsf{T}}\mathbf{z}_g$. This approach avoided explicit matrix inversion and improved numerical stability. Parameter combinations that produced nonfinite variances or a non-positive-definite covariance matrix were assigned a large objective-function value. Starting values for $\beta_0$ and $\beta_1$ were obtained from an ordinary least-squares regression fitted to all retained observations. To initialize the residual variance, separate ordinary least-squares regressions were fitted within groups, and their residual sums of squares were pooled across the corresponding residual degrees of freedom. Starting values for $\tau_a^2$ and $\tau_b^2$ were set to one half of the empirical variances among the separately estimated group intercepts and slopes, respectively. Each starting variance was constrained to exceed a small numerical floor equal to $10^{-10}$ times the larger of the total response variance and one. The negative log-likelihood was minimized using the Broyden–Fletcher–Goldfarb–Shanno quasi-Newton algorithm implemented by the `BFGS` method in the R function `optim`. No analytical gradient was supplied, so derivatives were evaluated numerically by `optim`. The maximum number of optimization iterations was 1,000 unless otherwise specified. The Hessian matrix was not requested because the fitted parameters were used to define the generative null distribution rather than to obtain Wald standard errors. Optimization was considered to have converged cleanly when `optim` returned convergence code zero and a finite negative log-likelihood. Nonzero convergence codes were retained as diagnostic warnings. The resulting estimates,

$$\widehat{\beta}_0, \quad \widehat{\beta}_1, \quad \widehat{\tau}_a^2, \quad \widehat{\tau}_b^2, \quad \widehat{\sigma}_e^2, \tag{14}$$

therefore define the maximum-likelihood member of the assumed linear Gaussian model family subject to zero latent intercept–slope covariance. Equivalently, this fitted model is the independent latent-parameter distribution that provides the greatest likelihood for the complete row-level observations after accounting for the observed predictor design and residual sampling variation. It is not fitted specifically to reproduce the observed correlation among separately estimated group coefficients. For the parametric bootstrap, latent group intercepts and slopes were independently sampled as

$$\alpha_g^* \sim \mathcal{N}\left(\widehat{\beta}_0, \widehat{\tau}_a^2\right), \tag{15}$$

$$\gamma_g^* \sim \mathcal{N}\left(\widehat{\beta}_1, \widehat{\tau}_b^2\right), \tag{16}$$

and simulated responses were generated at the original group-specific predictor values according to

$$y_{ig}^* = \alpha_g^* + \gamma_g^* x_{ig} + \varepsilon_{ig}^*, \qquad \varepsilon_{ig}^* \sim \mathcal{N}\left(0, \widehat{\sigma}_e^2\right). \tag{17}$$

Separate unconstrained ordinary least-squares reaction norms were then re-estimated from each simulated dataset. Although the latent intercepts and slopes were independent by construction, the recovered coefficients could be correlated because they were estimated from shared observations, finite samples, and the original predictor design. The resulting bootstrap distribution therefore represents the correlation or covariance expected among estimated reaction-norm parameters under the best-fitting latent-independence model.

### References


[1] Alex R. Gunderson. "Trade-offs between baseline thermal tolerance and thermal tolerance plasticity are much less common than it appears". In: *Global Change Biology* 29.13 (2023), pp. 3519–3524. DOI: 10.1111/gcb.16710. eprint: https://onlinelibrary.wiley.com/doi/pdf/10.1111/gcb.16710. URL: https://onlinelibrary.wiley.com/doi/abs/10.1111/gcb.16710.

[2] Sean W. Deery et al. "Heat hardening in a pair of Anolis lizards: constraints, dynamics and ecological consequences". In: *Journal of Experimental Biology* 224.7 (Apr. 2021), jeb240994. ISSN: 0022-0949. DOI: 10.1242/jeb.240994. eprint: https://journals.biologists.com/jeb/article-pdf/224/7/jeb240994/1832555/jeb240994.pdf. URL: 10.1242/jeb.240994.

[3] Rachael Morgan, Mette H. Finnøen, and Fredrik Jutfelt. "CT max is repeatable and doesn't reduce growth in zebrafish". In: *Scientific Reports* 8.1 (2018), p. 7099. DOI: 10.1038/s41598-018-25593-4. URL: 10.1038/s41598-018-25593-4.

[4] Anthony L. Gilbert and Donald B. Miles. "Antagonistic Responses of Exposure to Sublethal Temperatures: Adaptive Phenotypic Plasticity Coincides with a Reduction in Organismal Performance". In: *The American Naturalist* 194.3 (2019), pp. 344–355. DOI: 10.1086/704208. URL: 10.1086/704208.

[5] Belinda van Heerwaarden, Vanessa Kellermann, and Carla M. Sgrò. "Limited scope for plasticity to increase upper thermal limits". In: *Functional Ecology* 30.12 (2016), pp. 1947–1956. DOI: 10.1111/1365-2435.12687. URL: 10.1111/1365-2435.12687.

[6] Matthew C. Sasaki and Hans G. Dam. "Genetic differentiation underlies seasonal variation in thermal tolerance, body size, and plasticity in a short-lived copepod". In: *Ecology and Evolution* 10.21 (2020), pp. 12200–12210. DOI: 10.1002/ece3.6851. URL: 10.1002/ece3.6851.

[7] Matthew C. Sasaki and Hans G. Dam. "Negative relationship between thermal tolerance and plasticity in tolerance emerges during experimental evolution in a widespread marine invertebrate". In: *Evolutionary Applications* 14.8 (2021), pp. 2114–2123. DOI: 10.1111/eva.13270. URL: 10.1111/eva.13270.

[8] Andrew D. Nguyen et al. "Trade-Offs in Cold Resistance at the Northern Range Edge of the Common Woodland Ant Aphaenogaster picea (Formicidae)". In: *The American Naturalist* 194.6 (2019), E151–E163. DOI: 10.1086/705939. eprint: https://www.journals.uchicago.edu/doi/pdf/10.1086/705939. URL: https://www.journals.uchicago.edu/doi/abs/10.1086/705939.

[9] Giovanna Mottola et al. "Are you ready for the heat? Phenotypic plasticity versus adaptation of heat tolerance in three-spined stickleback". In: *Ecosphere* 13.4 (2022), e4015. DOI: 10.1002/ecs2.4015. eprint: https://esajournals.onlinelibrary.wiley.com/doi/pdf/10.1002/ecs2.4015. URL: https://esajournals.onlinelibrary.wiley.com/doi/abs/10.1002/ecs2.4015.

[10] Suegene Noh et al. "Seasonal variation in basal and plastic cold tolerance: Adaptation is influenced by both long- and short-term phenotypic plasticity". In: *Ecology and Evolution* 7.14 (2017), pp. 5248–5257. DOI: 10.1002/ece3.3112. eprint: https://onlinelibrary.wiley.com/doi/pdf/10.1002/ece3.3112. URL: https://onlinelibrary.wiley.com/doi/abs/10.1002/ece3.3112.

[11] Matthew C. Sasaki and Hans G. Dam. "Integrating patterns of thermal tolerance and phenotypic plasticity with population genetics to improve understanding of vulnerability to warming in a widespread copepod". In: *Global Change Biology* 25.12 (2019), pp. 4147–4164. DOI: 10.1111/gcb.14811. eprint: https://onlinelibrary.wiley.com/doi/pdf/10.1111/gcb.14811. URL: https://onlinelibrary.wiley.com/doi/abs/10.1111/gcb.14811.

[12] Erica O'Neill, Hannah E. Davis, and Heath A. MacMillan. "A lack of repeatability creates the illusion of a trade-off between basal and plastic cold tolerance". In: *Proceedings of the Royal Society B: Biological Sciences* 288.1964 (Dec. 2021), p. 20212121. ISSN: 0962-8452. DOI: 10.1098/rspb.2021.2121. eprint: https://royalsocietypublishing.org/rspb/article-pdf/doi/10.1098/rspb.2021.2121/623334/rspb.2021.2121.pdf. URL: 10.1098/rspb.2021.2121.

[13] Yimen G. Araya-Ajoy and Niels J. Dingemanse. "Repeatability, heritability, and age-dependence of seasonal plasticity in aggressiveness in a wild passerine bird". In: *Journal of Animal Ecology* (2017). DOI: 10.1111/1365-2656.12621.

[14] Jolle W. Jolles et al. "Personality, plasticity and predictability in sticklebacks: Bold fish are less plastic and more predictable than shy fish". In: *Animal Behaviour* 154 (2019), pp. 193–202. DOI: 10.1016/j.anbehav.2019.06.022. URL: 10.1016/j.anbehav.2019.06.022.

[15] Joram Soch et al. *The Book of Statistical Proofs (2025 Edition)*. Accessed: 2026-05-26. 2025. DOI: 10.5281/zenodo.18096132. URL: 10.5281/zenodo.18096132.
